# NaBC1 Acts as a Mechanosensitive Co-regulator of Fibronectin-binding Integrins Adhesion and Myoblast Polarization

**DOI:** 10.1101/2025.11.17.688253

**Authors:** Juan Gonzalez-Valdivieso, Rafael R. Castillo, Aleixandre Rodrigo-Navarro, Ana Rodriguez-Romano, Hayk Mnatsakanyan, Manuel Salmeron-Sanchez, Patricia Rico

## Abstract

Cell–matrix interactions are central to the regulation of cell mechanics, polarity, and signal transduction. In this study, we investigate the contribution of the borate transporter NaBC1 to myoblast adhesion dynamics and mechanotransductive responses. Using C2C12 myoblasts cultured on fibronectin-coated substrates, we show that NaBC1 activation rapidly reinforces cell–substrate attachment, leading to accelerated spreading and the establishment of polarized cell morphologies. These changes are associated with the assembly of enlarged focal adhesions and a marked slowdown of actin retrograde flow, consistent with increased force transmission across adhesion sites. NaBC1 stimulation also induces a coordinated increase in the expression of fibronectin-binding integrins, including α_5_β_1_ and α_v_β_3_, at both transcriptional and protein levels. Proximity ligation assays reveal an enhanced spatial association between NaBC1 and these integrins at the cell membrane, supporting the formation of cooperative adhesion complexes. In parallel, fluorescently labelled boron accumulates at focal adhesions and within intracellular compartments such as mitochondria, lysosomes, and the endoplasmic reticulum, suggesting a link between NaBC1-dependent adhesion signaling and subcellular organization. Importantly, the adhesive and mechanotransductive effects driven by NaBC1 are strictly dependent on fibronectin integrity and are lost on mutant fibronectins lacking RGD or synergy motifs, as well as on laminin-111 substrates that do not support molecular clutch engagement. Together, these findings identify NaBC1 as an integral component of fibronectin–integrin adhesion systems, contributing to the regulation of myoblast mechanics and polarity through cooperative interactions with fibronectin-binding integrins.

## 1. Introduction

Cell adhesion enables cells to establish physical and biochemical connections with their surrounding microenvironment, allowing them to withstand mechanical forces and coordinate tissue organization^1,2^. Through adhesion to the extracellular matrix (ECM), cells continuously sense mechanical cues such as tension, compression, and rigidity, which are translated into intracellular signals that regulate proliferation, migration, differentiation, and survival^3,4^. These mechanosensitive responses are essential for tissue development, maintenance, and regeneration^5–9^.

Cell–ECM interactions are primarily mediated by transmembrane adhesion receptors, among which integrins play a central role^10–13^. Integrins are heterodimeric receptors composed of α and β subunits that physically link ECM ligands to the actin cytoskeleton^10^. Upon engagement with the matrix, integrins cluster into focal adhesions (FAs), multiprotein complexes that include adaptor and signaling proteins such as talin, kindlin, vinculin, and paxillin^14–18^. These structures organize into a stratified adhesome that enables force transmission and mechanochemical signaling^19,20^. Through this molecular architecture, integrins regulate how cells adapt their cytoskeleton and behavior to ECM mechanical properties^10,21^.

The dynamic coupling between integrins and actin filaments is often described by the molecular clutch model, which explains how actin retrograde flow is modulated by substrate stiffness and adhesion strength^15–17^. When integrin–ECM bonds are weak, actin filaments slide rearward with minimal force transmission. In contrast, strong integrin engagement slows actin flow, promotes FA maturation, and increases intracellular tension^15–17^. This mechanism underlies essential cellular processes such as polarization, migration, and force generation.

In addition to integrins, several membrane proteins have emerged as modulators of mechanosensitive adhesion. Mechanically activated ion channels such as Piezo1 have been shown to cooperate with integrins, influencing FA signaling and cell–ECM communication^22,23^. More recently, the boron (B) transporter NaBC1 (*SLC4A11*), a Na⁺-coupled borate cotransporter essential for B homeostasis has been identified as a membrane protein capable of influencing mechanotransductive responses^24–27^.

Mutations in *SLC4A11* are associated with corneal endothelial dystrophies^25^, highlighting its physiological relevance. Beyond its transport function, NaBC1 has been implicated in diverse adhesion-dependent cellular processes, including osteogenic differentiation^26^, inhibition of adipogenesis^27^, promotion of angiogenesis^28^, adhesion-driven osteogenesis^29^, myogenic differentiation and muscle regeneration through crosstalk with growth factor receptors (GFRs)^30,31^. These studies suggest that NaBC1 participates in signaling pathways that extend beyond ion transport.

Previous work demonstrated that NaBC1 cooperates with fibronectin (FN)-binding integrins to regulate myoblast responses to substrate stiffness and intracellular tension^24^. However, whether NaBC1 also contributes to early cell polarization, a process tightly linked to FA maturation and cytoskeletal asymmetry, remains unknown. Cell polarization establishes directional organization of actin structures and adhesion sites and precedes migration, fusion, and differentiation in muscle cells^32,33^.

Here, we investigate the role of NaBC1 in early myoblast polarization and adhesion dynamics. Using C2C12 myoblasts cultured on FN-functionalized substrates, we show that NaBC1 activation enhances FA formation, reduces actin retrograde flow, and promotes rapid cell spreading and polarization. These effects are accompanied by transcriptional and protein-level upregulation of FN-binding integrins (α_5_β_1_ and α_v_β_3_) and their spatial colocalization with NaBC1 at the cell membrane. Importantly, by employing recombinant FN variants lacking RGD or synergy motifs, we demonstrate that NaBC1-mediated adhesion reinforcement depends on the structural integrity of FN-integrin binding sites. Together, these findings identify NaBC1 as a cooperative component of the FN-integrin mechanosensitive machinery that regulates adhesion strength, cytoskeletal tension, and polarization in myoblasts. This mechanism provides new insights into how ion transporters integrate with ECM recognition to control muscle cell behavior and may be relevant for pathological conditions involving defective cell-ECM interactions.

## 2. Results and Discussion

To investigate muscle cell adhesion, C2C12 murine myoblasts were employed as a widely accepted *in vitro* model of skeletal muscle cells^34^. Building on previous studies showing that the coordinated activation of FN-binding integrins (α_5_β_1_ and α_v_β_3_) together with NaBC1 enhances myogenic differentiation in C2C12 cells^30^, this work focuses on dissecting the specific contribution of NaBC1 to adhesion-driven cellular responses. Earlier reports demonstrated that NaBC1-mediated adhesion states regulate lineage commitment in mesenchymal stem cells (MSC)^29^, accelerate myotube formation and muscle repair *in* vivo^31^ and establish NaBC1 as a mechanosensitive regulator of myoblast adhesion through its interaction with FN-binding integrins and intracellular tension^26^. Here, we extend these findings by specifically analyzing early adhesion behavior. To selectively stimulate NaBC1 and FN-binding integrins, soluble borax salt (B) and FN were used, respectively.

As an initial step, the potential cytotoxicity of B was assessed in C2C12 myoblasts cultured under proliferative conditions in serum-containing growth medium. Cell proliferation was evaluated after 24 and 48 h of exposure to increasing B concentrations. As shown in **Supplementary Figure 1**, B concentrations above 52 mM (20 mg mL⁻¹) significantly impaired cell viability, whereas lower concentrations did not adversely affect proliferation. Based on these results, a concentration of 0.59 mM (corresponding to 0.2 mg mL⁻¹) was selected for all subsequent experiments. At this concentration, cell viability remained above 95%, and no cytotoxic effects were detected. In addition, this dose was chosen in agreement with previous studies reporting its biological activity in different cell types, including C2C12 myoblasts^28–31^.

### 2.1. Active NaBC1 promotes C2C12 cell spreading and polarization under potassium-depleted conditions

To further examine whether NaBC1 activation can drive adhesion-dependent responses, we evaluated early cell behavior under potassium-depleted conditions (**Figure 1**). Potassium (K^+^) is known to play a critical role in the maturation of cell adhesion structures^35^, and its depletion has been reported to impair the formation of mature focal adhesions (FA) and prevent cell polarization despite allowing initial attachment^36^. Based on this, intracellular K⁺ levels were reduced by culturing C2C12 myoblasts in K⁺-free medium, following previously established protocols^36^. Representative images of cell morphology under the different experimental conditions are shown in Fig. 1A, while quantitative analysis of cell and nuclear parameters is presented in Fig. 1B. Cells were cultured on FN-coated glass substrates in complete medium (CM) or potassium-depleted medium (–K), in the presence or absence of borax (B), and analyzed at 0.5, 1, and 1.5 h. In CM, cells adhered rapidly and initiated spreading within the first 30 minutes. After 1.5 h, they displayed an elongated and polarized morphology, accompanied by the formation of FA at the cell periphery and along protrusions. In contrast, cells cultured under potassium depletion exhibited a markedly different phenotype. Although attachment still occurred, cells remained rounded and spread radially, showing a circumferential actin organization and lacking the anisotropic architecture associated with FA maturation and stress fiber formation (Fig. 1A). Quantitatively, potassium depletion resulted in a significant increase in cell circularity (approximately threefold) and a pronounced reduction in spreading area. In addition, these cells retained a refractile nuclear appearance (Fig. 1A, arrows), indicative of incomplete spreading.

**Figure 1.**
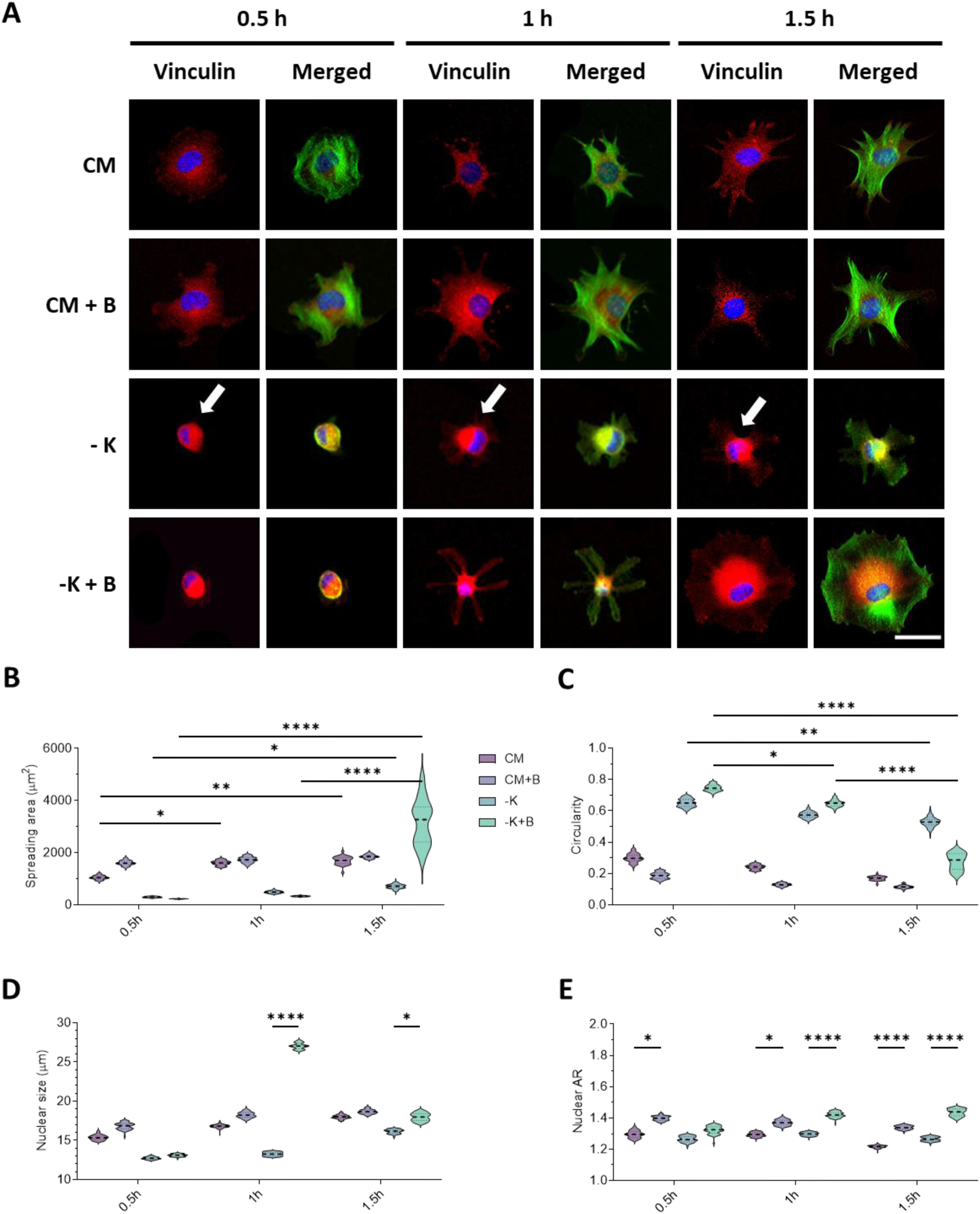
Active-NaBC1 induces C2C12 cell polarization and spreading after potassium (K^+^) depletion. A: Representative images of C2C12 myoblasts seeded on FN-functionalized glasses and cultured for 0.5 h, 1 h and 1.5 h with complete medium (CM) or K^+^-depleted medium (-K) with 0.59 mM borax (+B). Samples without B are considered as a control of each condition. Arrows: perinuclear refractility characteristic of partially spread cells. Green: actin; Red: vinculin; Blue: DAPI. Scale bar: 25 μm. B-E: Quantified cell spreading area (B), cell circularity (C), nuclear size (D) and nuclear aspect ratio (E) of C2C12 myoblasts, cultured as described in panel A. n > 50 imaged cells/condition from three different biological replicas. All data are presented as Mean ± SD. Statistical significance was determined using two-way ANOVA (Tukey’s post hoc test) for multiple comparisons. *p ≤ 0.05, **p ≤ 0.01, ****p ≤ 0.0001

Strikingly, the addition of B promoted a substantial recovery of cell spreading and polarization under both CM and –K conditions. Cells treated with B showed reduced circularity and a marked increase in projected area, reaching values up to ∼3150 µm² (Fig. 1B). This effect was accompanied by progressive changes in nuclear morphology, including increased nuclear size and aspect ratio over time, suggesting enhanced cytoskeletal tension and mechanical coupling between the cytoskeleton and the nucleus. Consistent with this interpretation, cells exposed to B displayed more prominent actin stress fibers (Fig. 1A), indicative of an adhesion state capable of sustaining intracellular tension

### 2.2. Active NaBC1 enhances early focal adhesion assembly and strengthens cell-substrate attachment

Following the analysis of cell morphology, we next examined the formation of FAs, as these structures provide the physical linkage between the extracellular matrix (ECM) and the actin cytoskeleton and are essential for the regulation of adhesion strength and cell behavior^37^. FA assembly is a stepwise process initiated by integrin engagement with the matrix, which triggers the recruitment of adaptor and signaling proteins at the cell membrane^10,21^. Early adhesion complexes are characterized by the accumulation of proteins such as focal adhesion kinase (FAK), paxillin, and talin, which subsequently promote the incorporation of additional components, including vinculin and zyxin, leading to adhesion maturation and stabilization^38,39,40^. The dynamic regulation of these structures is tightly coupled to actin organization and is modulated by signaling pathways such as those governed by Rho GTPases^39^.

To quantify FA formation, vinculin staining was used as a marker (**Figure 2**A). C2C12 myoblasts were seeded at low density (5 x 10^3^ cells cm^-2^) on FN-coated glass substrates in serum-free conditions to selectively promote α_5_β_1_ and α_v_β_3_ integrin engagement while minimizing cell-cell interactions. Morphological parameters together with FA number and size were quantified over time in control and B-treated cells (Figs. 2B-F). NaBC1 activation induced a clear increase in the number of FAs at early time points. After 1 and 1.5 h of culture, cells treated with B exhibited approximately 100 and 93 adhesions per cell, respectively, compared to 52 and 65 adhesions in control conditions. In addition to this increase in number, FA morphology was also altered. Adhesions in NaBC1-activated cells displayed larger dimensions, with structures reaching up to ∼12 µm, in contrast to the smaller, punctate adhesions observed in control cells. These differences were already evident after 0.5 h, where B-treated cells presented more than 150 adhesions per cell together with an increase in adhesion area. The enlargement of FAs is typically associated with enhanced mechanical coupling between the cell and the substrate, reflecting stronger adhesion forces, whereas smaller and more transient adhesions are linked to higher cell motility^41^. In this context, the observed increase in both FA number and size upon NaBC1 activation suggests the establishment of a reinforced adhesion state at early stages, consistent with the enhanced spreading and polarization described above.

**Figure 2.**
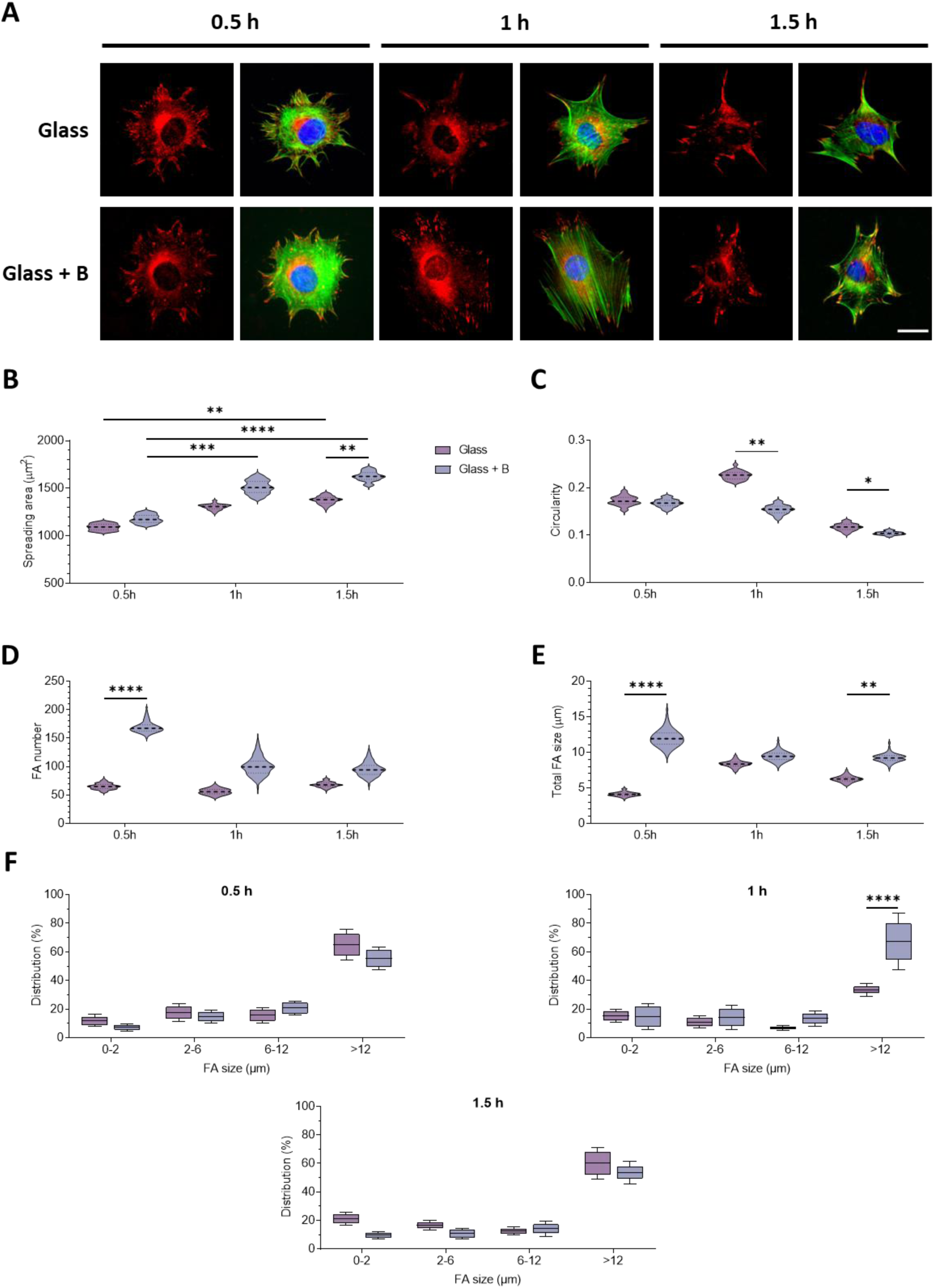
Active-NaBC1 induces focal adhesion formation and C2C12 cell adhesion at early stages. A: Representative images of C2C12 myoblasts seeded on FN-functionalized glasses and treated with 0.59 mM B for 0.5, 1 and 1.5 h. Green: actin; Red: vinculin; Blue: DAPI. Scale bar: 25 μm. B-C: Quantified morphological cell parameters indicative of cell spreading area (B) and cell circularity (C) of C2C12 myoblasts, cultured as described in panel A. n > 50 imaged cells/condition from three different biological replicas. D-F: Quantified number of focal adhesions (D), FA size (E) and FA size distribution (F) in C2C12 myoblasts, cultured as described in panel A. n > 50 imaged cells/condition from three different biological replicas. All data are presented as Mean ± SD. Statistical significance was determined using two-way ANOVA (Tukey’s post hoc test) for multiple comparisons. *p ≤ 0.05, **p ≤ 0.01, ***p ≤ 0.001, ****p ≤ 0.0001

### 2.3. NaBC1 activation regulates actin cytoskeleton dynamics and force transmission

The dynamics of the actin cytoskeleton play a central role in cell migration and adhesion, with retrograde actin flow representing a key feature of these processes^42^. Depending on cell type and migration mode, actin flow can be localized at protrusive regions such as lamellipodia or extend throughout the entire cell body^43,44^. This rearward movement of actin filaments, relative to the substrate, results from the balance between actin polymerization at the leading edge and contractile forces generated by actomyosin activity^45^. The coupling of this actin movement to the ECM through integrin-based adhesions enables the generation of traction forces required for cell spreading and motility^45^. The efficiency of this coupling is commonly described by the molecular clutch model, in which the engagement between the actin cytoskeleton and integrin-mediated adhesions determines whether actin flow is dissipated or converted into mechanical work^14–16^. Weak interactions between the cytoskeleton and the matrix lead to rapid actin flow and limited force transmission, whereas stronger adhesion promotes slower actin movement and increased loading of FA components, including talin unfolding and vinculin recruitment^14–16,46^. This behavior is also influenced by substrate properties, with stiffer environments favoring more stable adhesion structures and enhanced cytoskeletal organization.

To assess whether NaBC1 activation influences actin dynamics, retrograde actin flow was quantified in live C2C12 myoblasts (**Figure 3**A-B). Based on the increased adhesion observed in previous sections, we hypothesized that NaBC1 stimulation would enhance cytoskeletal coupling and consequently reduce actin flow. Consistent with this, cells treated with B (0.59 mM) displayed a significant decrease in retrograde actin velocity compared to untreated cells, with values of 5.07 nm s⁻¹ and 7.78 nm s⁻¹, respectively. To further explore the dependence of this effect on the extracellular matrix context, actin flow was also measured in cells cultured on laminin-111-coated substrates. Laminin has been reported to alter cellular sensitivity to substrate mechanics^47^. Under these conditions, NaBC1 activation did not produce detectable changes in actin flow, which remained at approximately 10 nm s⁻¹. These values are comparable to those previously reported for cells cultured on rigid polyacrylamide substrates with similar mechanical properties^24^. Taken together, these results indicate that NaBC1 activation modulates actin cytoskeleton dynamics in a matrix-dependent manner. The reduction in retrograde actin flow observed on FN, combined with the enhanced adhesion and polarization described above (Figs. 1 and 2), supports the idea that NaBC1 promotes a more effective mechanical coupling between the actin cytoskeleton and integrin-based adhesions. In this context, NaBC1 appears to contribute to the engagement of the molecular clutch, facilitating force transmission to the extracellular matrix and reinforcing adhesion-dependent mechanotransduction in myoblasts.

**Figure 3.**
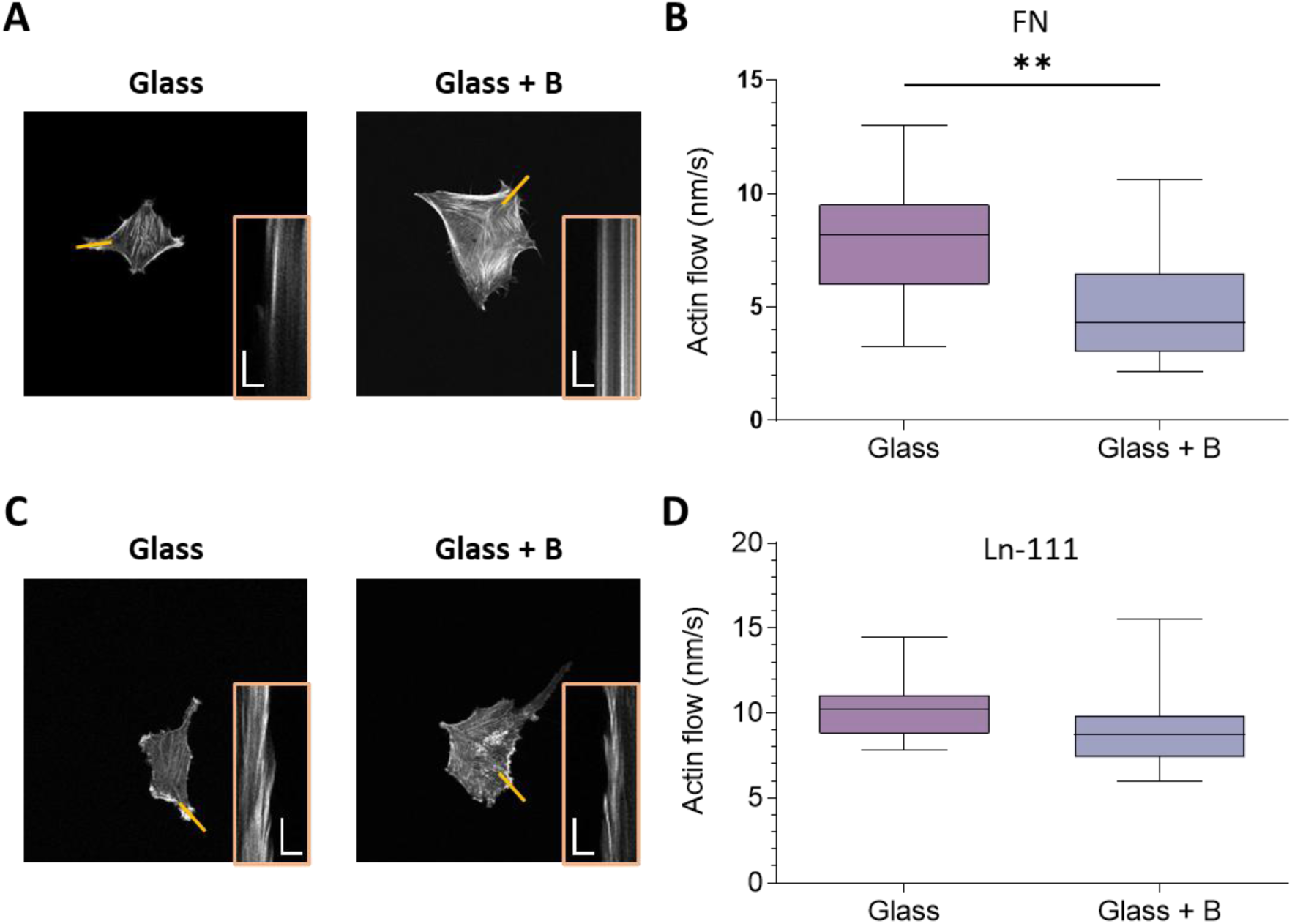
Stimulation of NaBC1 modulates actin cytoskeleton dynamics on FN. A: Representative images of C2C12 myoblasts transfected with LifeAct-GFP, seeded on FN-functionalized glasses and treated with 0.59 mM B for 1.5 h. Yellow lines: location where kymographs were computed, on average. Insets: kymographs showing the movement of actin features. The spatial scale bar in the inset is 5 μm, whereas the temporal scale bar is 1 minute. B: Quantified actin retrograde flow in C2C12 myoblasts, cultured as described in panel A. n = 5 cells with at least 5 different flow areas per cell. C: Representative images of C2C12 myoblasts transfected with LifeAct-GFP, seeded on laminin-111-functionalized glasses and treated with 0.59 mM B for 1.5 h. Yellow lines: location where kymographs were computed, on average. Insets: kymographs showing the movement of actin features. The spatial scale bar in the inset is 5 μm, whereas the temporal scale bar is 1 minute. D: Quantified actin retrograde flow in C2C12 myoblasts, cultured as described in panel C. n = 5 cells with at least 5 different flow areas per cell. All data are presented as Mean ± SD. Statistical significance was determined using t-test for pairwise comparisons. **p ≤ 0.01

### 2.4. Active-NaBC1 promotes FN-binding integrin expression and membrane receptor association

Integrin expression is dynamically regulated by multiple factors, including ECM composition, intracellular signaling pathways, and mechanical cues such as substrate stiffness and tensile forces^48,12,49,50^. In particular, FN-binding integrins (α_5_β_1_ and α_v_β_3_) play a central role in controlling cell adhesion, migration, and survival^10,15^. Emerging evidence also indicates that ion-channels and transporters can influence integrin expression and function through signaling crosstalk mechanisms^51–53^. Although the specific molecular pathways remain incompletely defined, several studies have linked ion fluxes and channel activity to integrin regulation via pathways involving calcium signaling, Rho GTPases, and FAK activation^54–56.57,58,53,52^. These observations suggest that membrane transport systems may participate in the modulation of integrin-mediated adhesion.

Multiple studies have previously reported ion-channels and ion-transporters physically coupling together with integrins to produce functional clusters at the cell membrane^59,60^. These clusters trigger reciprocal signaling and integrin-channel crosstalk, wherein cell adhesion may trigger channel activation and channel engagement can control cell adhesion^61,62^. The location of ion-channels and transporters in the plasma membrane may be influenced by integrins and these multiprotein clusters. In fact, ion-channels can occasionally use conformational coupling for downstream signaling^63^. As is the case with several ion-channels that pair with integrin β_1_ and stimulate its expression^64^, the channel protein frequently feeds back by regulating integrin activation and/or expression. FN-binding integrins, such α_5_β_1_ and α_v_β_3_, have a regulatory effect on the healing processes and greatly enhance muscle stability^65,66^, as well as to modulate cell adhesion, differentiation, and migration^67^.

Based on this framework, we investigated whether NaBC1 activation affects the expression of FN-binding integrins in C2C12 myoblasts. Quantitative analysis revealed a rapid upregulation of the genes encoding α_5_, α_v_, β_1_, and β_3_ integrin subunits following NaBC1 stimulation (**Fig. 4**A–B). This effect was detected as early as 0.5 h and persisted over time (1 and 1.5 h). Among these, α_v_ and β_3_ exhibited the most pronounced increase, reaching approximately 2.4-fold and 2-fold higher expression levels, respectively, compared to unstimulated cells on glass substrates. The early onset of this transcriptional response correlates with the increased number and size of FAs observed at similar time points (Fig. 2), suggesting a coordinated regulation of adhesion assembly and integrin expression.

**Figure 4.**
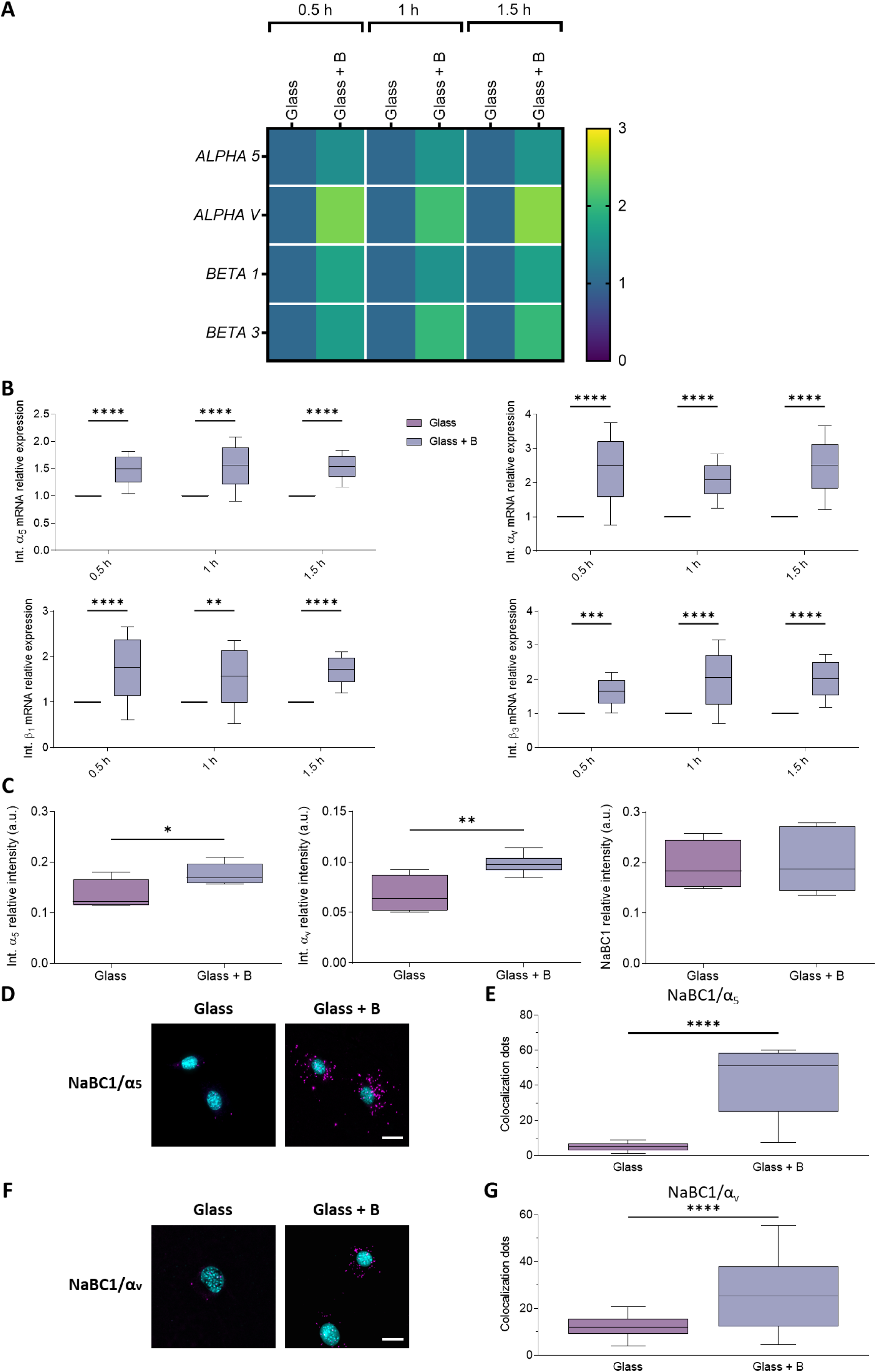
Active-NaBC1 upregulates FN-binding integrins expression and NaBC1-integrin colocalization. A: Heat map of quantitative real-time PCR analysis of relative mRNA expression of cell adhesion-related genes (ALPHA 5, ALPHA V, BETA 1, BETA 3 Integrins) in C2C12 myoblasts seeded on FN-functionalized glasses and treated with 0.59 mM B for 0.5, 1, and 1.5 h, as measured by qPCR. B: Quantified relative mRNA expression of cell adhesion-related genes showed in panel A. n = 3 biological replicates with 3 technical replicates. C: Quantified expression of Integrin α_5_, Integrin α_v_ and NaBC1 by In-Cell Western in C2C12 myoblasts seeded on FN-functionalized glasses and treated with 0.59 mM B. n > 3 biological replicates with 3 technical replicates. D and F: Representative images showing the colocalization dots of NaBC1/α_5_ (D) and NaBC1/α_v_ (F) in C2C12 myoblasts seeded on FN-functionalized glasses and treated with 0.59 mM B for 3 h. Magenta: colocalization dots; Cyan: DAPI. Scale bar: 50 μm. E and G: Quantified number of colocalization dots of NaBC1/α_5_ (E) and NaBC1/α_v_ (G). n = 30 cells from 3 different biological replicates. All data are presented as Mean ± SD. Statistical significance was determined using one-way ANOVA, two-way ANOVA (Tukey’s post hoc test) or t-tests for multiple or pairwise comparisons, respectively. *p ≤ 0.05, **p ≤ 0.01, ****p ≤ 0.0001

To determine whether these changes were also reflected at the protein level, α_5_ and α_v_ integrins were quantified using In-Cell Western assays. As shown in Figure 4C, NaBC1 activation resulted in a consistent increase in integrin protein levels, in agreement with the transcriptional data. In contrast, NaBC1 protein expression itself did not show significant variation upon B treatment, consistent with previous observations^31^.

Given the upregulation of FN-binding integrins, we next evaluated the spatial relationship between NaBC1 and α_5_/α_v_ integrins using a proximity ligation assay (DUOLINK® PLA) (Fig. 4D–G and **Supplementary Fig. S2**). This approach allows the detection of protein-protein proximity within a range of approximately 40 nm. NaBC1 stimulation led to a marked increase in the number of PLA signals, indicating enhanced colocalization with both α_5_ (Fig. 4D–E) and α_v_ (Fig. 4F–G) integrins after 3 h of culture. These results support the formation of NaBC1–integrin complexes at the cell membrane.

Notably, a significant increase in NaBC1-integrin proximity was already observed after 30 minutes of stimulation (Fig. S2), a time point at which cells have not yet produced or reorganized their own FN matrix. This early response indicates that NaBC1 associates with integrins engaged with pre-adsorbed FN, rather than relying on cell-derived ECM. Therefore, NaBC1-integrin coupling appears to be an initial event in the adhesion process. Taken together, these findings demonstrate that NaBC1 activation not only enhances the expression of FN-binding integrins but also promotes their spatial association at the cell membrane. This coordinated response likely contributes to the stabilization of adhesion sites (Fig. 2), the acquisition of polarized morphology (Fig. 1), and the strengthening of actin-integrin coupling reflected in reduced actin retrograde flow (Fig. 3). In this context, NaBC1 emerges as a cooperative component of integrin-based adhesion complexes, linking membrane transport activity with mechanochemical signaling at the cell-ECM interface.

### 2.5. NaBC1 activation enhances myotube formation and modulates integrin remodeling during myogenic differentiation

After demonstrating that NaBC1 activation increases the expression of FN-binding integrins in myoblasts, we next assessed whether these effects were maintained during myogenic differentiation into multinucleated myotubes (**Figure 5**A). In agreement with previous observations^30,31^, stimulation with soluble B significantly enhanced myotube formation. Quantitative analysis revealed an increase in the fusion index together with a marked enlargement of myotube width, reaching values of 21.09 µm compared to 5.75 µm in untreated controls (Fig. 5B). In addition to these morphological changes, cytoskeletal organization was also affected. Actin staining intensity within myotubes was approximately doubled in the B-treated condition (Fig. 5C-D), indicating the formation of a more developed and structured contractile network. These features are consistent with an advanced stage of myotube maturation.

**Figure 5.**
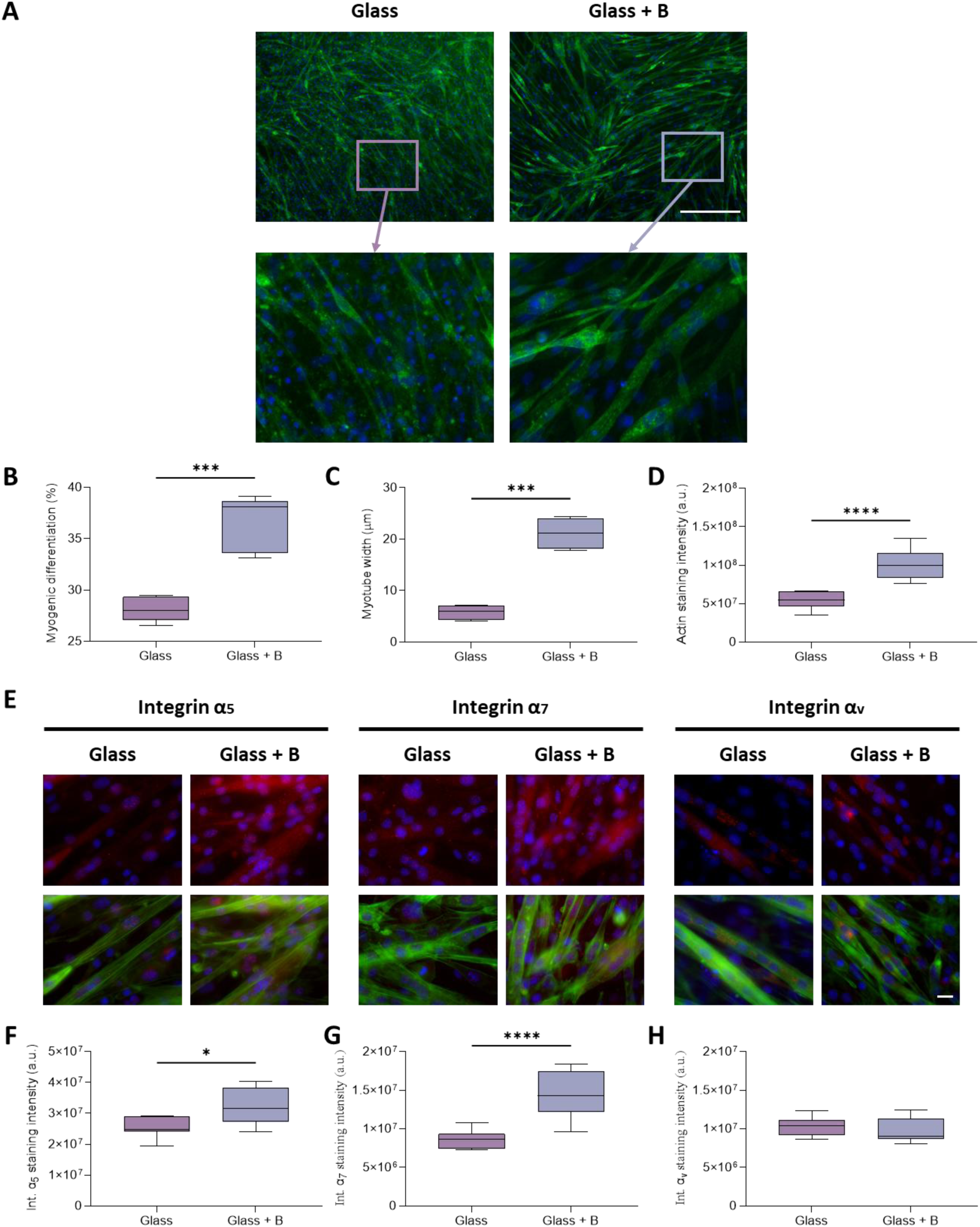
Active-NaBC1 upregulates integrins protein expression in myotubes. A: Representative images of myogenic differentiation in C2C12 myoblasts seeded on FN-functionalized glasses and treated with 0.59 mM B for 96 h. Green: actin; Blue: DAPI. Scale bar: 100 μm. Zoom areas are indicated. B and C: Quantified myotube differentiation and myotube width, cultured as described in panel A. n > 20 imaged cells/condition from three different biological replicas. D: Quantified actin intensity in myotubes, cultured as described in panel A. n > 20 imaged cells/condition from three different biological replicas. E: Representative images of myogenic differentiation in C2C12 myoblasts, cultured as described in panel A. Green: actin; Blue: DAPI. Scale bar: 100 μm. F-H: Quantified α_5_, α_7_, and α_v_ integrins protein expression in myotubes, cultured as described in panel A. n > 50 imaged cells/condition from three different biological replicas. All data are presented as Mean ± SD. Statistical significance was determined using t-test for pairwise comparisons. *p ≤ 0.05, ***p ≤ 0.001, ****p ≤ 0.0001

NaBC1 activation also altered integrin expression profiles during differentiation. Protein-level quantification showed a significant increase in α_5_ and α_7_ integrin levels in B-treated myotubes relative to controls (Fig. 5E–H), while α_v_ integrin exhibited a similar upward trend. These findings indicate that NaBC1 not only enhances morphological differentiation but also modulates adhesion receptor composition during myogenesis.

Integrin expression is known to be tightly regulated throughout muscle differentiation^68–70^. Proliferating myoblasts predominantly express α_5_β_1_ integrin, whereas differentiated myotubes progressively shift toward α_7_β_1_, the principal laminin (LN) receptor in mature skeletal muscle^69,71^. Interestingly, despite the expected reduction of α_5_ expression during myotube formation, our results show sustained levels of this integrin under NaBC1 stimulation. This observation points to a potential cooperative role for FN-binding integrins during differentiation, possibly contributing to enhanced cell-matrix interactions required for efficient fusion and cytoskeletal organization. Previous studies have shown that α_5_β_1_ supports early adhesion and fusion events, while α_7_β_1_ is essential for the structural stability and function of mature muscle fibers^70,72^.

Altogether, these results indicate that NaBC1 regulates integrin remodeling during myogenesis by promoting α_7_ expression while maintaining α_5_ levels. This dual modulation may reinforce adhesion to both FN and LN-rich environments, thereby supporting more efficient myotube formation and potentially contributing to improved muscle regeneration.

### 2.6. Boron localizes at adhesion sites and intracellular organelles upon NaBC1 activation, linking adhesion to cellular bioenergetics

To investigate the intracellular distribution of boron following NaBC1 activation, C2C12 myoblasts were incubated with fluorescein-labelled boron (FITC-B). Confocal imaging revealed a pronounced accumulation of FITC-B at the cell periphery (**Fig. 6**A), suggesting enrichment at regions associated with cell-substrate interactions. This observation is consistent with the previously described spatial association between NaBC1 and FN-binding integrins (Fig. 4-D, E, F, G), raising the possibility that B uptake occurs in proximity to FA sites. To evaluate this, colocalization between FITC-B and vinculin was quantified in cells cultured on FN- or LN-111-coated substrates (Fig. 6B). The degree of spatial overlap was assessed using the Manders Overlap Coefficient (MOC), a widely used metric for fluorescence colocalization ranging from 0 (no overlap) to 1 (complete overlap)^73^. Cells cultured on FN exhibited higher MOC values (0.89) compared to LN-111 conditions (0.80) (Fig. 6B–C), indicating a stronger association of FITC-B with FAs in FN environments. These results support the idea that B internalization is spatially linked to adhesion structures and is influenced by ECM context.

**Figure 6.**
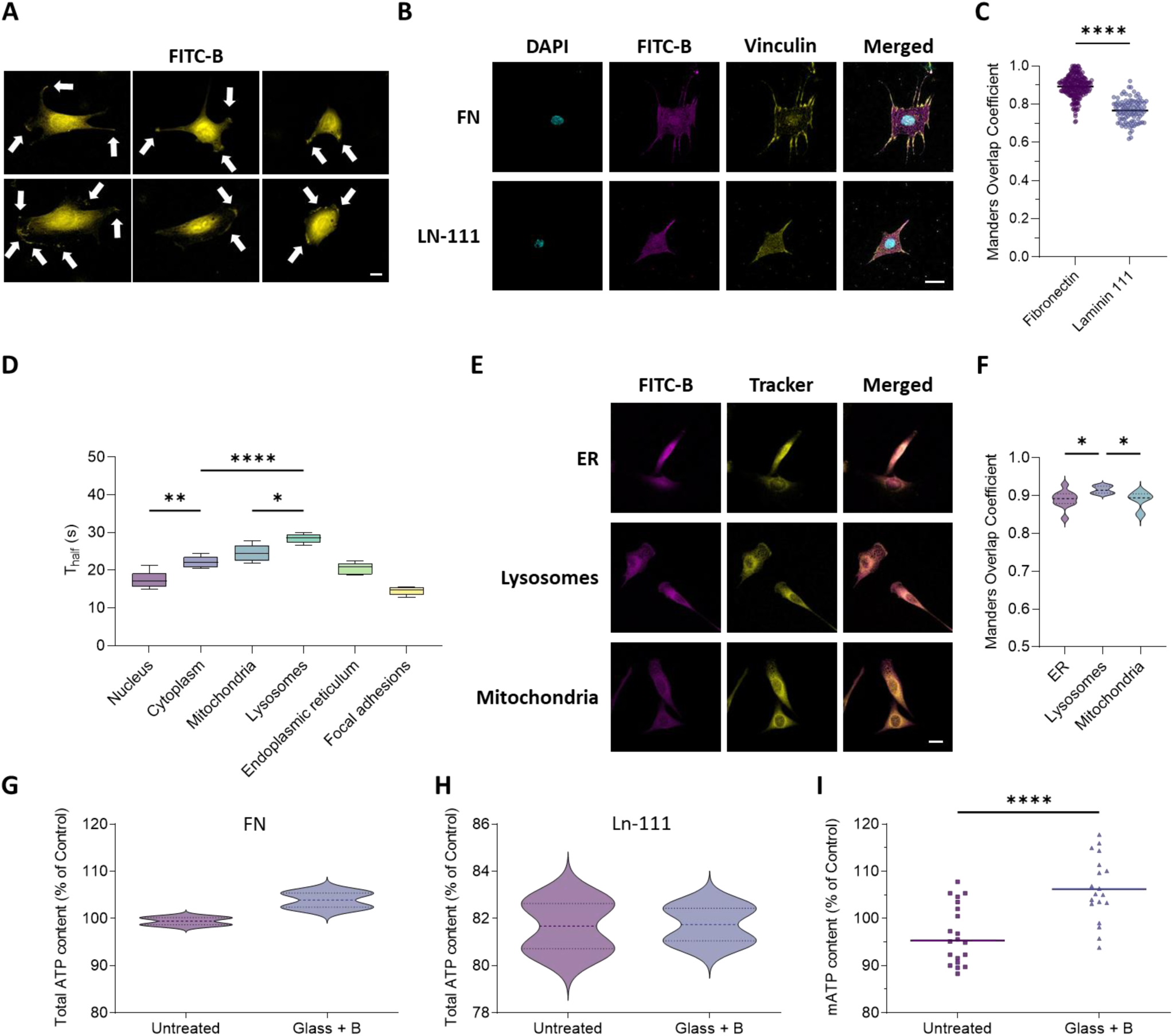
Fluorescent labelled FITC-B colocalizes within FAs and accumulates in intracellular organelles. A: Representative images of the subcellular localization of FITC-B in C2C12 myoblasts seeded on FN-functionalized glasses and treated with FITC-labelled Boron (FITC-B) for 1 h. Yellow: FITC-B. Arrows: clumps in cell edges. Scale bar: 20 μm. B: Representative images of FITC-B and focal adhesions colocalization in C2C12 myoblasts seeded on glass substrates functionalized with FN or laminin-111 and treated with FITC-B for 1 h. Magenta: FITC-B; Yellow: vinculin; Cyan: DAPI. Scale bar: 20 μm. n = 20 cells from 3 different biological replicates. C: Quantified FITC-B and focal adhesion colocalization in C2C12 cells seeded on glass substrates functionalized with FN or laminin-111 and treated with FITC-B for 1 h, as scored using the Manders Overlap Coefficient (MOC). D: Quantified half-life times FITC-labelled B in C2C12 myoblasts, cultured as described in panel A. n = 10 cells from 3 different biological replicates. E: Representative images of the colocalization of FITC-B and live trackers in C2C12 myoblasts seeded on FN-functionalized glasses and treated with FITC-B for 1 h. Magenta: FITC-B. Yellow: live trackers for ER (ER-Tracker), lysosomes (LysoTracker) and mitochondria (MitoTracker). Scale bar: 20 μm. F: Quantified FITC-B and live trackers colocalization in C2C12 myoblasts, cultured as described in panel E. n: 3 biological replicates with 3 technical replicates. G: Quantified total ATP content in C2C12 myoblasts seeded on FN-functionalized glasses and treated with 0.59 mM B. n: 3 biological replicates with 3 technical replicates. H: Quantified total ATP content in C2C12 myoblasts seeded on laminin-111-functionalized glasses and treated with 0.59 mM B. n: 3 biological replicates with 3 technical replicates. I: Quantified mitochondrial ATP content in C2C12 myoblasts, cultured as described in panel G. n: 3 biological replicates with 3 technical replicates. All data are presented as Mean ± SD. Statistical significance was determined using one-way ANOVA (Tukey’s post hoc test) or t-tests for multiple or pairwise comparisons, respectively. *p ≤ 0.05, ****p ≤ 0.0001

To further characterize the dynamics of B within cells, Fluorescence Recovery After Photobleaching (FRAP) experiments were performed (**Supplementary Fig. S3**). FITC-B displayed a relatively rapid turnover within FAs, with a half-life of approximately 15 s (Fig. 6D), suggesting transient residence or continuous exchange at these sites. In contrast, longer retention times were observed in intracellular organelles, including lysosomes (∼29 s) and mitochondria (∼25 s), indicating preferential accumulation within these compartments. Co-staining with organelle-specific markers confirmed high levels of colocalization between FITC-B and lysosomes, mitochondria, and the endoplasmic reticulum, with coefficients ranging from 0.88 to 0.91 (Fig. 6E–F).

Given the prominent localization of B within mitochondria, we next examined whether NaBC1 activation influences cellular energy metabolism. Although the relationship between ECM cues and metabolic activity is still being elucidated^74–76^, cytoskeletal remodeling is known to impose a substantial energetic demand, accounting for a significant fraction of ATP consumption in various cell types^77,78^. Therefore, total ATP and mitochondrial ATP (mATP) levels were measured in C2C12 myoblasts cultured under the different conditions (Fig. 6G–I).

Myoblasts stimulated with B showed a modest but consistent increase in both total ATP (104 % vs. 98 % in controls) and mitochondrial ATP levels (107 % vs. 95 %) when cultured on FN-coated substrates (Fig. 6G and I). In contrast, no significant changes in ATP content were observed in cells cultured on LN-111 (Fig. 6H), further supporting the matrix-dependent nature of NaBC1-mediated responses. Collectively, these findings extend the functional role of NaBC1 beyond its involvement in cell adhesion, revealing a connection between adhesion-related signaling and intracellular metabolic activity. The enrichment of B at FAs suggests its participation in integrin-associated processes, while its accumulation in mitochondria and other organelles points to a broader role in cellular homeostasis. The increase in mitochondrial ATP production upon NaBC1 activation indicates that this transporter may contribute to the metabolic adaptation required to sustain cytoskeletal remodeling and adhesion strengthening. In this context, NaBC1 emerges as a link between mechanotransduction and bioenergetic regulation in myoblast^79,80^.

### 2.7. NaBC1-integrin complexes preferentially activate Akt signaling

Although integrins, particularly β1-containing complexes, are known to be essential for myoblast fusion both *in vitro* and *in vivo*^81^, the downstream signaling pathways that mediate this process remain incompletely understood. To identify the intracellular mechanisms associated with NaBC1 activation, we first examined focal adhesion kinase (FAK), a well-established mediator of integrin signaling at FAs^38,82^. Phosphorylation of FAK at Tyr397 (pFAK) is commonly associated with adhesion-dependent signaling during myogenic differentiation^83^. However, analysis of FAK activation revealed no significant differences in pFAK levels or in the pFAK/FAK ratio following NaBC1 stimulation (**Supplementary Fig. S4**A–B). These findings suggest that the enhanced adhesion and differentiation responses observed are not primarily mediated through canonical FAK-dependent pathways, in line with previous reports linking NaBC1 activation to contractility-related signaling downstream of FAK^28,29^.

FN-binding integrins and the Akt pathway are both involved in regulating various aspects of cell behavior, including cell adhesion, migration, proliferation, and survival^10,21,84,85^. The phosphatidylinositol 3-kinase (PI3K)/Akt pathway is a major signaling pathway that regulates many cellular processes, including cell growth, survival, and metabolism^86^. Activation of the Akt pathway is often triggered by receptor tyrosine kinases or other signaling molecules, which activate PI3K and lead to the activation of Akt through a series of phosphorylation events^87,88^. Previous studies suggest that the engagement of α_5_β_1_ and α_v_β_3_ integrins with FN induces Akt activation thereby promoting cell survival, migration, and proliferation in different cell types^85,89^. The interplay between FN-binding integrins and the Akt pathway is complex, multifaceted and involves bidirectional signaling and regulation between these two pathways. Conversely, the Akt pathway can also regulate the expression and activity of FN-binding integrins. Thus, activation of the Akt pathway has been shown to increase the expression of α_5_β_1_ integrin in breast cancer cells, and to promote the formation of FAs in response to FN^85^. Moreover, Akt activation has been associated with increased integrin expression and FA assembly through downstream effectors such as integrin-linked kinase (ILK)^90,91^. Previous studies from our group also reported that B treatment enhances Akt expression and phosphorylation^24,29^. On the other hand, it is widely accepted that integrins influence Akt pathway and, indeed, the PI3K/Akt signaling pathway has been previously reported to undergo adhesion-dependent activation^92^. Considering the significant upregulation of α_5_ and α_v_ integrins upon NaBC1 activation (Fig. 4A–C), we investigated whether Akt signaling was modulated under these conditions. Western blot and In-Cell Western analyses showed a clear increase in Akt phosphorylation at Ser473 in B-treated cells compared to controls (Fig. S4C–E), indicating activation of the Akt pathway.

The differential activation of Akt in the absence of changes in pFAK suggests that NaBC1/FN-integrin clustering selectively engage specific signaling adhesion pathways. While integrin–FAK signaling is typically associated with FA turnover and migration processes, our data suggest that NaBC1 preferentially couples with FN-binding integrins to potentiate Akt-driven pathways, which are more directly linked to cell survival, cytoskeletal organization, and metabolic regulation. In the context of myogenesis, Akt is a key regulator of myoblast fusion and hypertrophic growth, supporting its relevance in the differentiation process.

Taken together, these results indicate that NaBC1 activation promotes a signaling shift toward Akt-dependent pathways, rather than classical FAK-mediated signaling. This selective coupling provides a mechanistic framework to explain the enhanced adhesion (Figs. 1-2), cytoskeletal reinforcement (Fig. 3), myotube formation (Fig. 5), and metabolic adaptation (Fig. 6) observed in this study, linking NaBC1-integrin clustering to downstream pathways that coordinate muscle cell maturation and function.

### 2.8. NaBC1 activation reduces myoblast migration and fibronectin matrix remodeling

Cell migration and extracellular matrix (ECM) remodeling are tightly coordinated processes involved in tissue morphogenesis, regeneration, and pathological events such as fibrosis and tumor invasion^93–95^. Migrating cells continuously interact with and mechanically remodel their surrounding matrix through traction forces generated by the actomyosin cytoskeleton. These forces promote ECM deformation, fiber alignment, FA turnover, and ECM reorganization, all of which contribute to directional migration and tissue restructuring^93,95^. In turn, ECM architecture and mechanical properties influence cell polarity, adhesion dynamics, and migratory behavior. Matrix stiffness, ligand composition, and ECM organization are therefore critical regulators of cell motility^94,95.96^. Cells tend to migrate more efficiently on matrices that are similar in stiffness to the surrounding tissue, and changes in matrix stiffness can alter the migration speed and direction of cells^96^.

During skeletal muscle regeneration, satellite cells (SCs) become activated and give rise to proliferating myoblasts that migrate toward the damaged area before differentiating and fusing into multinucleated myofibers^97.98,99^. Because these events require coordinated regulation of adhesion and motility, we next examined whether NaBC1 activation influences FN remodeling and myoblast migration (**Figure 7**). Quantification of FN reorganization revealed that NaBC1 activation significantly reduced the extent of FN fibril reorganization. The remodeled FN area decreased from approximately 1350 µm² in control conditions to ∼730 µm² in B-treated cells (Fig. 7A–B). Consistent with this observation, time-lapse migration assays demonstrated that NaBC1 stimulation markedly impaired cell motility. B-treated myoblasts migrated shorter distances and at lower velocities than untreated cells, with migration distances decreasing from ∼460 µm to ∼170 µm and migration speed from 0.003 µm s⁻¹ to 0.0017 µm s⁻¹ (Fig. 7C–E).

**Figure 7.**
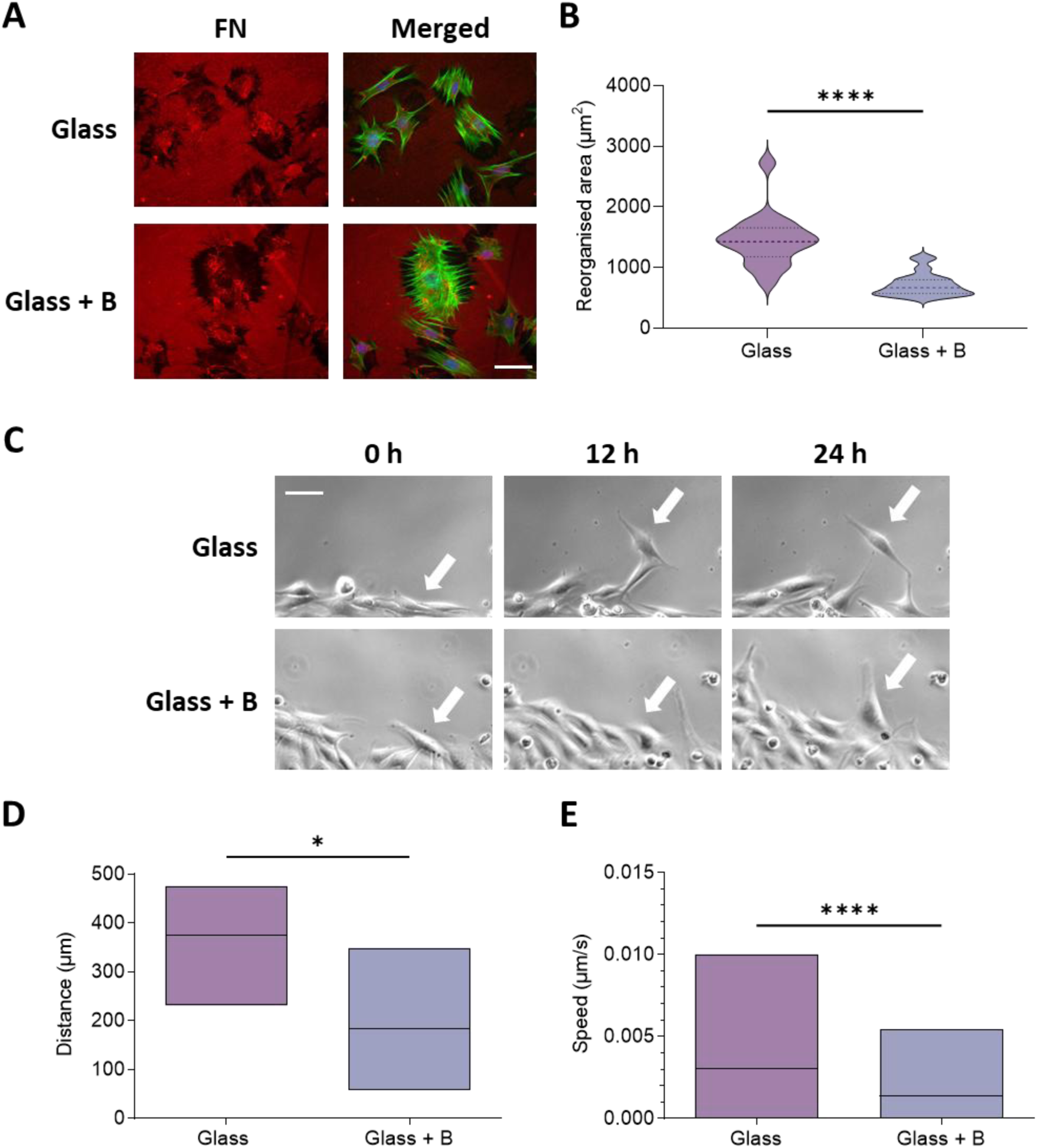
Active NaBC1 reduces myoblast migration and FN reorganization. A: Representative images of the matrix reorganization in C2C12 myoblasts seeded on FN-functionalized glasses and treated with 0.59 mM B. Green: actin; Red: fibronectin; Blue: DAPI. Dark area around each cell was quantified to establish the reorganization area. Scale bar: 50 μm. B: Quantified matrix reorganization in C2C12 myoblasts, cultured as described in panel A. n = 10 images from 3 different biological replicas. C: Representative images of C2C12 myoblasts, cultured as described in panel A, after 0 h, 12 h and 24 h. Arrows: position of a selected migrating cell over time in different conditions. Scale bar: 50 μm. D: Quantified total distance covered by tracked cells from time-lapse microscopy experiments. n = 6 different tracked cells per condition and 139 measurements per cell for a total of 24 h. E: Quantified speed average covered by tracked cells from time-lapse microscopy experiments. n = 6 different tracked cells per condition and 139 measurements per cell for a total of 24 h. All data are presented as Mean ± SD. Statistical significance was determined using t-test for pairwise comparisons. *p ≤ 0.05, ****p ≤ 0.0001

These findings are in agreement with the enhanced adhesive phenotype described throughout this study. NaBC1 activation promotes FA maturation, increases cytoskeletal organization, and reduces retrograde actin flow, all of which are associated with stronger mechanical coupling between the cell and the ECM. According to the molecular clutch model^14–16,37^, increased force transmission through integrin-based adhesions stabilizes FAs and limits their turnover, thereby reducing the dynamic detachment-reattachment cycles required for efficient migration and ECM remodeling.

In this context, the reduced migratory behavior observed upon NaBC1 activation likely reflects a shift from a motile to a more mechanically anchored state. Because efficient FN remodeling depends on continuous traction-mediated reorganization of the matrix, the stabilization of integrin-ECM interactions induced by NaBC1 would also be expected to limit ECM deformation. Thus, NaBC1 appears to regulate the balance between adhesion and motility by favoring stable adhesion over migratory dynamics. From a myogenic perspective, these results suggest that NaBC1 may exert stage-dependent functions during muscle regeneration. Whereas early myoblast migration and ECM remodeling are necessary to populate damaged regions, the subsequent acquisition of a more adhesive and less migratory phenotype may facilitate cell fusion, cytoskeletal stabilization, and maturation into multinucleated myotubes.

### 2.9. NaBC1 cooperation with FN-binding integrins depends on RGD and synergy sites

We previously showed that the activation of the NaBC1 transporter markedly influences cell spreading and adhesion through specific crosstalk with FN-binding integrins^24,31^. To further investigate how NaBC1 interacts with FN-binding integrins and how this cooperation affects adhesion and mechanotransduction, C2C12 myoblasts were cultured on recombinant FNs containing specific mutations in their integrin-binding domains (**Figure 8 and Figure Supplementary 5**). The wild-type FN (WT-FN) fragment (FNIII_9–10_) contains both the RGD sequence and the adjacent synergy site (PHSRN), which together mediate high-affinity binding to α_5_β_1_ integrins. The RGD sequence is essential for integrin binding, particularly for α_5_β_1_ and α_v_β_3_, while the synergy site reinforces α_5_β_1_- FN interactions under mechanical tension, stabilizing FAs and promoting downstream signaling^100,101^.

**Figure 8.**
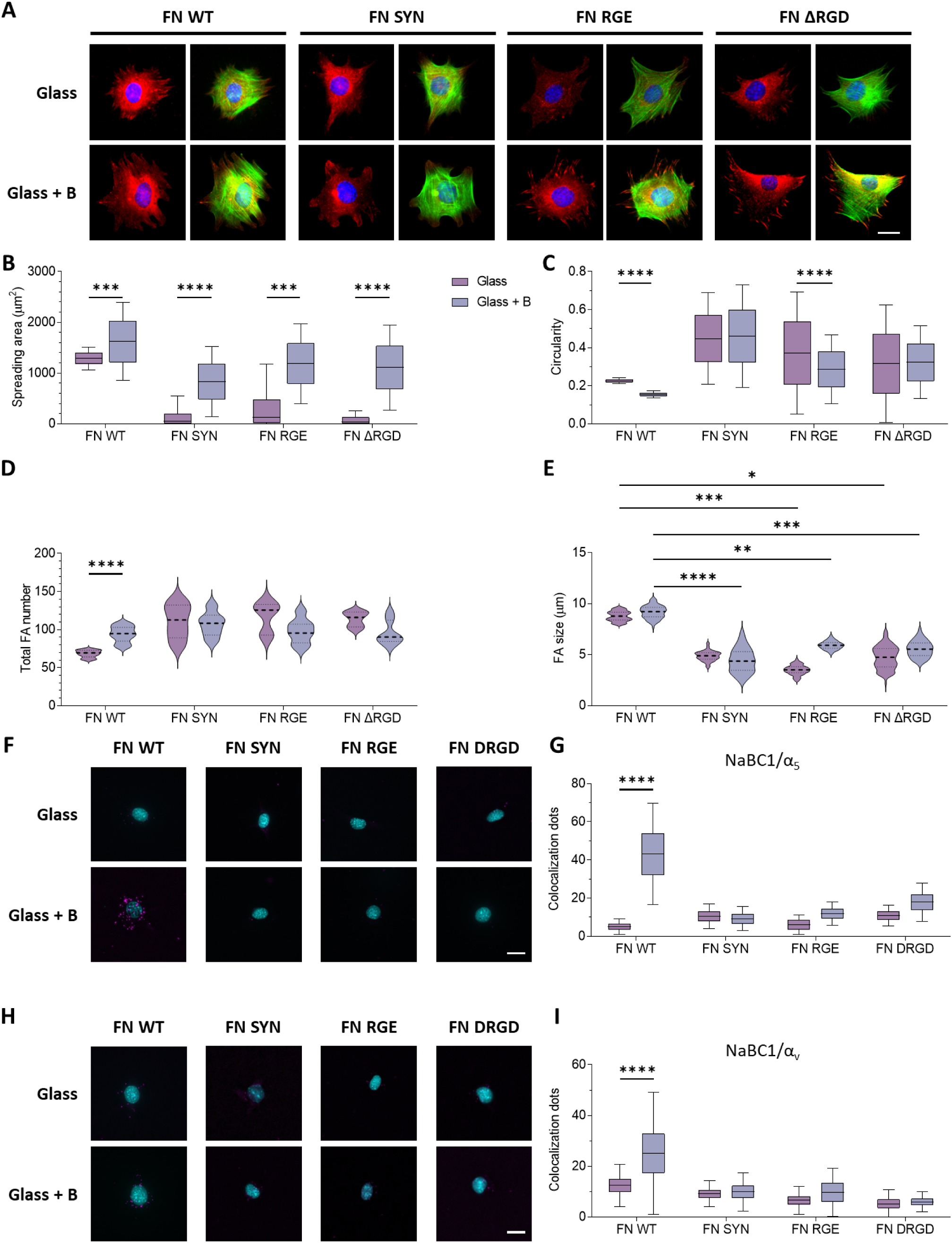
The effects of active-NaBC1 on cell adhesion is dependent on RGD and synergy sites of fibronectin. A: Representative images of C2C12 myoblasts seeded on glass substrates functionalized with WT or mutant FN forms and treated with 0.59 mM B for 1.5 h. Green: actin; Red: vinculin; Blue: DAPI. Scale bar: 25 μm. B: Quantified cell spreading area of C2C12 myoblasts, cultured as described in panel A. n > 50 imaged cells/condition from three different biological replicas. C: Quantified cell circularity of C2C12 myoblasts, cultured as described in panel A. n > 50 imaged cells/condition from three different biological replicas. D: Quantified number of FAs in C2C12 myoblasts, cultured as described in panel A. n > 50 imaged cells/condition from three different biological replicas. E: Quantified FA size in C2C12 myoblasts, cultured as described in panel A. n > 50 imaged cells/condition from three different biological replicas. F and H: Representative images showing the colocalization dots of NaBC1/α_5_ (F) and NaBC1/α_v_ (H) in C2C12 myoblasts seeded on glass substrates functionalized with WT or mutant FN forms for 3 h and treated with 0.59 mM B. Magenta: colocalization dots; Cyan: DAPI. Scale bar: 50 μm. G and I: Quantified number of colocalization dots of NaBC1/α_5_ (G) and NaBC1/α_v_ (I). n= 30 cells from 3 different biological replicates. All data are presented as Mean ± SD. Statistical significance was determined using two- way ANOVA (Tukey’s post hoc test) for multiple comparisons. *p ≤ 0.05, **p ≤ 0.01, ***p ≤ 0.001, ****p ≤ 0.0001

Mutant FN variants lacking the RGD motif (FNΔRGD), the synergy site (FNΔsyn), or containing a point mutation in which the aspartic acid in RGD is replaced by glutamic acid (FN-RGE) were used to dissect the contribution of each motif. The FN-RGE mutant serves as a functional negative control, as this substitution disrupts integrin recognition while preserving the overall structure of the FN fragment.

Cells seeded on WT-FN exhibited strong adhesion and well-organized actin stress fibers, which were further enhanced upon NaBC1 activation with soluble B (0.59 mM B). In contrast, cells on FNΔRGD, FNΔsyn or FN-RGE displayed reduced spreading area and increased circularity compared to WT-FN (Fig. 8A, B, C). Moreover, myoblasts on mutant FNs presented fewer and smaller FAs upon NaBC1 activation (Fig. 8D, E), indicating that NaBC1 stimulation failed to rescue adhesion in any of these mutant substrates, confirming that the NaBC1-mediated reinforcement of adhesion depends on functional FN-integrin interactions through both the RGD and synergy motifs.

Interestingly, the effect of interaction between NaBC1 and FN-binding integrins showed a strong change regarding the distribution of the FAs sizes when myoblasts were cultured onto mutant FNs (**Figure Supplementary S5**). Thus, the activation of NaBC1 transporter induced the formation of larger FAs on WT-FN (60% of FAs >12 µm) compared to cells seeded on mutant FN forms, where around 80% of FAs were smaller than 2 µm.

These findings reveal that NaBC1 cooperates with α_5_β_1_ integrins in a manner that depends on the integrity of the RGD and synergy sites. The FN-RGE control confirms that NaBC1 cannot engage or signal through FN in the absence of functional integrin binding, ruling out nonspecific mechanisms. This supports a model in which NaBC1 may participate in the same adhesion complex as α_5_β_1_, reinforcing FN-integrin bonds under tension and amplifying FA maturation and mechanotransductive signaling^12,100^.

Colocalization analysis (Fig. 8F-I) further supported these findings, showing strong colocalization of NaBC1 with α_5_ and α_v_ integrins on WT-FN, that was not reproduced on mutant FNΔRGD, FNΔsyn, FN-RGE, suggesting that both motifs are also essential for NaBC1-integrin spatial colocalization and highlights the requirement for both chemical and mechanical integrity of the FN-integrin interface. Together, these findings indicate that NaBC1 acts as a cooperative component of the α_5_β_1_-FN mechanosensitive complex, stabilizing adhesion structures.

## 3. Conclusions

This work identifies the boron transporter NaBC1 as a regulator of cell adhesion and mechanotransduction in myoblasts, acting in close cooperation with FN-binding integrins. We show that NaBC1 activation by soluble B enhances cell adhesion strength, actin cytoskeletal organization, and intracellular tension, promoting a more adhesive and polarized phenotype while reducing cell migration and FN reorganization.

By using recombinant FN variants lacking specific integrin-binding motifs (ΔRGD, Δsyn, and RGE), we further demonstrate that NaBC1-mediated adhesion and signaling depend on the integrity of the RGD and synergy sites. The absence of these motifs abolishes the adhesion-driven state and reduces NaBC1-integrin colocalization, confirming a structural and functional interdependence between NaBC1 and FN-binding integrins.

Altogether, our results define NaBC1 as a multifunctional mechanosensor that integrates B signaling with ECM recognition and intracellular force generation. Through its cooperative action with FN-binding integrins at FN-binding domains, NaBC1 strengthens cell-matrix adhesion, modulates cytoskeletal tension, and regulates cell polarization. This integrative mechanism establishes NaBC1 as a key modulator of myoblast behavior and a potential target for controlling adhesion-driven processes in muscle regeneration and biomaterial design.

## 4. Experimental section

### 4.1. Material substrates

Cleaned glass cover slips were used as 2D control substrates for *in vitro* experiments. Sodium Tetraborate Decahydrate Borax 10 Mol (hereafter borax -B-) (Na_2_B_4_O_7_·10H_2_O, Borax España S.A) was dissolved in the culture medium at 0.59 mM.

All the glasses were functionalized with human plasma FN (Sigma-Aldrich) unless otherwise specified. After sterilizing under UV for 30 minutes, the glasses were coated with 20 µg mL^-1^ FN solution in Dulbecco’s Phosphate Saline Buffer (DPBS) at room temperature for 1 h.

### 4.2. C2C12 myoblasts culture

Murine C2C12 myoblasts (Sigma-Aldrich) were cultured in Dulbecco’s Modified Eagle Medium (DMEM, Invitrogen) with high glucose content (4.5 g L^-1^), supplemented with 20 % Fetal Bovine Serum (FBS, Invitrogen) and 1 % antibiotics (P/S; 10,000 units mL^-1^ of penicillin and 10,000 µg mL^-1^ streptomycin per 100 mL of media, Thermofisher) in humidified atmosphere at 37° C and 5 % CO_2_. Cells were sub-cultured before reaching confluence (approximately every 2 days).

### 4.3. Cytotoxicity assay

MTS quantitative assay (The CellTiter 96 Aqueous One Solution Cell Proliferation Assay, Promega) was performed to assess cytocompatibility of borax with C2C12 cells and to establish the maximum working concentrations. 2 x 10^4^ cells cm^-2^ were seeded onto a p-24 multi-well plate and metabolic activity was measured after 48 h of incubation with different concentrations of borax (0.2, 0.6, 3, 6, 10 and 20 mg mL^-1^) in cell culture media. Cells were then incubated for 3 h with MTS (tetrazolium salt) at 37° C and the formation of formazan was followed by measuring absorbance at 490 nm. All measurements were performed in triplicate.

### 4.4. Cell adhesion experiments

For cell adhesion experiments with potassium depletion, C2C12 myoblasts were seeded at low density of 5 x 10^3^ cells cm^-2^ on FN-functionalized glasses. Intracellular potassium was depleted by incubation in K^+^-free medium for 30 minutes, as previously described^36^. C2C12 myoblasts were then cultured in complete medium (CM - 140 mM NaCl, 50 mM Hepes, 1 mM CaCl_2_, 0,5 mM MgCl_2,_ 3 mM KCl) or potassium depleted medium (-K - 140 mM NaCl, 50 mM Hepes, 1 mM CaCl_2_, 0,5 mM MgCl_2_). Cells were supplemented with borax 0.59 mM as required. After 0.5 h, 1 h and 1.5 h of culture, cells were washed in DPBS (Gibco) and fixed in 4 % formaldehyde solution (Sigma-Aldrich) for 30 minutes at 4° C. Three biological replicas were evaluated.

For cell adhesion experiments, C2C12 cells were seeded at low density of 5 x 10^3^ cells cm^-2^ on FN-functionalized glasses. Cells were cultured in DMEM with high glucose content, 1 % P/S antibiotics in absence of FBS. Cells were also supplemented with borax 0.59 mM as required. After 0.5, 1 and 1.5 h of culture, cells were washed in DPBS and fixed in 4 % formaldehyde solution at 4° C for 30 minutes. Three biological replicas were evaluated.

### 4.5. Immunofluorescence assays

Cells from adhesion or matrix reorganization assays were rinsed with DPBS and permeabilized with DPBS/0.5 % Triton x-100 at room temperature for 5 minutes, blocked in 2% BSA/DPBS at 37° C for 1 h, and then incubated with primary antibodies in blocking solution at 4° C overnight. After incubation with the primary antibodies, samples were rinsed twice in DPBS/0.1 % Triton X-100 and incubated with the secondary antibody and/or BODIPY FL phallacidin (Invitrogen, 1:100) at room temperature for 1 h. Finally, samples were washed twice in DPBS/0.1 % Triton X-100 and mounted with Vectashield containing DAPI (Vector Laboratories). Imaging was performed under an epifluorescence microscope (Nikon Eclipse 80i).

Monoclonal primary antibody against vinculin for focal adhesion detection (Sigma-Aldrich, 1:400) and Cy3 conjugated secondary antibody (Jackson Immunoresearch, 1:200) were employed for cell adhesion assays.

### 4.6. Actin flow

C2C12 myoblasts were transfected with the LifeAct-GFP plasmid (Ibidi) using the Neon transfection system (ThermoFisher Scientific) following manufacturer’s protocol. The parameters used to achieve cell transfection were 1650 V, 10 ms, 3 pulses, and 5 µg of DNA. Transfected cells were cultured for 24 h before seeding.

Cells were seeded at 1×10^4^ cells cm^-2^ on functionalized glasses and allowed to adhere for 24 h. Imaging was performed using a ZEISS LSM900 confocal microscope with a 40x oil-immersion objective and ZEN software. Images were acquired at 488 nm at a frame rate of 1 image every 2 seconds for 4 minutes in total. 5 cells with at least 5 different flow areas per cell were measured.

### 4.7. Gene expression

Total RNA from C2C12 myoblasts cultured for 0.5 h, 1 h and 1.5 h was extracted using RNeasy micro kit (Qiagen). RNA quantity and quality was measured with a NanoDrop 1000 (ThermoFisher Scientific). 500 ng of RNA were reverse transcribed using the Superscript III reverse transcriptase (Invitrogen) and oligo dT primer (Invitrogen). Real-time qPCR was performed using SYBR select master mix in a 7500 Real Time PCR system (Applied Biosystems). At least three technical and biological replicas were measured. Sequences from the GenBank database were used to design the primers for amplification: Integrin α_5_ (NM_010577.3, Forward: 5’-GGA CGG AGT CAG TGT GCT G-3’, Reverse: 5’-GAA TCC GGG AGC CTT TGC TG-3’), Integrin β_1_ (XM_011248315.1, Forward: 5’-CAT CCC AAT TGT AGC AGG CG-3’, Reverse: 5’-CGT GTC CCA CTT GGC ATT CAT-3’), Integrin α_v_ (NM_008402.2, Forward: 5’-CAC CAG CAG TCA GAG ATG GA-3’, Reverse: 5’-GAA CAA TAG GCC CAA CGT CT-3’), Integrin β_3_ (NM_016780.2, Forward: 5’-GGA ACG GGA CTT TTG AGT GT-3’, Reverse: 5’-ATG GCA GAC ACA CTG GCC AC-3’). β-actin (NM_007393.3, Forward: 5’-TTC TAC AAT GAG CTG CGT GTG-3’, Reverse: 5’-GGG GTG TTG AAG GTC TCA AA-3’) was used as housekeeping gene. The analysis was performed by using the comparative Ct method based on the fractional cycle number at which fluorescence passed the threshold (Ct values). Sample values were normalized to the threshold value of β-actin (housekeeping gene): Δ*C*_*T*_ = *C*_*T*_(*experimental genes*) − *C*_*T*_ (βactin). ΔΔ*C*_*T*_ = Δ*C*_*T*_(*experimental sample*) − Δ*C*_*T*_(*control sample*). mRNA expression was calculated by the following equation: *fold c*ℎ*ange* = 2^−ΔΔ*CT*^

### 4.8. Intracellular signaling

In-Cell western was used to determine the protein levels of α_5_ and α_v_ integrins, NaBC1 transporter and pAkt/Akt ratio. 1 x 10^4^ C2C12 myoblasts cells cm^-2^ were seeded onto FN-functionalized glasses and treated with B (0.59 mM) for 1.5 h at 37° C and 5 % CO_2_. Cells were fixed using fixative buffer (10 mL formaldehyde, 90 mL PBS, 2 g sucrose) at 37° C for 15 minutes and then permeabilized with cold methanol at 40° C for 5 minutes. Samples were then blocked with 0.5 % blocking buffer (non-fat dry milk powder in 0.1 % PBST buffer) at RT for 2 hours followed by 3 washes with 0.1 % PBST for 10 minutes. After blocking, samples were incubated with primary antibodies diluted in blocking buffer at 4° C overnight. The primary antibodies used were: anti-NaBC1 (1:200, ABCAM), anti-integrin α_5_ (1:500, ABCAM), anti-integrin α_v_ (1:500 ABCAM) and anti-Akt/pAkt (1:500, Invitrogen). After 3 washes with 0.1 % PBST buffer for 10 minutes, samples were incubated with 1:800 diluted infrared-labeled secondary antibody IRDye 800CW (LI-COR) and 1:500 diluted CellTag 700 Stain (LI-COR) at RT for 1 hour, followed by 5 washes with 0.1% PBST for 10 minutes. Finally, samples were dried overnight at room temperature. Infrared signal was detected using an Odyssey infrared imaging system. Four biological replicas were analyzed.

Quantification of pFAK/FAK ratio was performed by western blot. Protein was extracted using RIPA buffer supplemented with protease inhibitor cocktail tablets (ROCHE) followed by protein separation in 10 % SDS-PAGE electrophoresis. Proteins were transferred to PVDF (GE Healthcare) membranes using Trans-blot Semi-dry transfer system (BIO-RAD). Then, membranes were blocked with 5 % BSA (Sigma-Aldrich) in TBS 0.1 % Tween 20 (Sigma-Aldrich) for 1 h and subsequently incubated with primary antibodies anti FAK (0.5 µg mL^-1^, Sigma-Aldrich), anti pFAK (1:500, Sigma-Aldrich) and anti α-tubulin as a loading control (1:5.000, ABCAM) diluted in TBS 0.1 % Tween 20 and 3 % BSA at 4° C overnight. After incubation, membranes were washed and incubated with HRP linked secondary antibodies (anti mouse or anti rabbit 1:10.000, GE Healthcare) at room temperature for 1 h. Protein bands were detected by chemiluminiscence using ECL-Plus (ThermoFisher) and visualized by Fujifilm Las-3000 imager device.

### 4.9. Colocalization experiments

Colocalization assays of NaBC1 and integrins were performed using DUOLINK^®^ PLA system (Sigma-Aldrich) following manufacturer instructions. The primary antibodies used were anti-NaBC1 (1:200, ABCAM), anti-integrin α_5_ (1:500, ABCAM) and anti-integrin α_v_ (1:500, ABCAM). At least 20 individual cells per condition were imaged using a Nikon Eclipse 80i epifluorescence microscope.

### 4.10. Chemical procedure for the preparation of Fluorescein-labelled B

Reactions were carried out using our previously reported procedure^24^. Briefly, solvents were purchased from Scharlab, while 4-(aminomethyl)phenylboronic acid pinacol ester hydrochloride, HCl in methanol (1.25 M), triethylamine, and sodium metaperiodate from Merck while the Fluorescein isothiocyanate (95% isomer mixture) from ABCR. TLC analyses were performed on silica gel supported onto aluminum foils. ^1^H NMR spectra were recorded on a Bruker Avance 400 using deuterated solvents (Merck), while FTIR and MS analyses recorded on a Perkin Elmer Spectrum two ATR spectrometer and an Agilent 6210 time-of-flight LC/MS respectively.

Target compound, 4-(fluorescein)thioureidomethylphenylboronic acid, was synthesized in two steps. First, the pinacol ester was cleaved oxidatively using NaIO₄ (240 mg, 1.1 mmol) and the boronic ester (250 mg, 0.93 mmol) in THF/water/HCl at room temperature for 4 h. After acid-finished work-up, the intermediate compound was obtained as a white solid (145 mg, 83%). The resulting boronic acid was then coupled with FITC (19.5 mg, 0.05 mmol) in EtOH basified with triethylamine (0.01 M in EtOH, 5 mL). The mixture was stirred for 2 h in the dark and purified by flash chromatography (EtOAc), yielding a bright orange solid (26 mg, 96%). All characterizations were coherent with previously reported data^24^.

### 4.11. Subcellular localization

1×10^4^ C2C12 myoblasts cm^-2^ were seeded on FN-functionalized glasses. After 24 h, cells were incubated with FITC-B at 37°C for 1 h. After washing with DPBS, cells were incubated with 75 nM LysoTracker^®^ Red DND-99 (Invitrogen), 100 nM MitoTracker^®^ Red CMXRos (Invitrogen) or 1 µM ER-tracker^®^ Red (Invitrogen) at 37° C for 1 h. 20 cells from 3 different biological replicates were imaged with a Zeiss LSM980 confocal microscope equipped with an incubator to maintain the temperature and humidity constant (37° C and 5 % CO_2_).

### 4.12. Fluorescence recovery after photobleaching (FRAP)

1×10^4^ C2C12 myoblasts cm^-2^ were seeded on FN-functionalized glasses. After 24 h, cells were treated with FITC-B for 1 h. Then, samples were washed with DPBS three times and imaged with a Zeiss LSM980 confocal microscope. 100 % fluorescence in the indicated areas (namely, cytoplasm, endoplasmic reticulum, focal adhesions, lysosomes, mitochondria and nucleus) was bleached for 2 ms with the 488 laser. Fluorescence recovery after bleaching was measured for 4 minutes. 10 cells from 3 different biological replicates were measured.

### 4.13. Total ATP content

1×10^4^ C2C12 myoblasts cm^-2^ were seeded on FN-functionalized glasses and allowed to adhere for 24 h. Cells were washed twice with PBS, lysed with 0.5 % Triton-X100 in PBS and samples were centrifuged at 11,000 × g for 3 minutes. 5 µl of total lysate were used to determine total ATP content using the ATP Determination Kit (ThermoFisher Scientific) following the manufacturer’s instructions. Luminescence at 560 nm was monitored using a Multiskan FC microplate reader (ThermoScientific). 3 biological replicates with 3 technical replicates were measured.

### 4.14. Mitochondrial ATP content

Mitochondrial ATP content was assessed with the BioTracker^TM^ ATP-Red dye (Millipore), as previously described^24^. 1×10^4^ C2C12 myoblasts cm^-2^ were seeded on FN-functionalized glasses. After 24 h, cells were incubated with 200 nM MTG for 1 h. Then, samples were washed twice with PBS and incubated for 15 minutes with 5 µM BioTracker^TM^ ATP-Red dye at 37° C and 5 % CO^2^. Then, the cells were washed twice with medium and fresh medium was added. Samples were imaged using a ZEISS LSM900 confocal microscope. At least 10 cells from 3 different biological replicates were measured.

### 4.15. Matrix reorganization and migration studies

5 x 10^3^ C2C12 myoblasts cm^-2^ were seeded on FN-functionalized glasses in the presence of 10 % FBS to allow cells to reorganize the FN. After 3 h, cells were washed with DPBS (Gibco) and fixed in 4 % formaldehyde solution (Sigma-Aldrich) for 30 minutes at 4° C. Following the same procedure explained in the immunofluorescence assay section, samples were incubated with anti-FN primary antibody (1:400, Sigma-Aldrich), Cy3-conjugated anti-mouse secondary antibody (1:200, Jackson Immunoresearch) and BODIPY FL phallacidin (1:100, Invitrogen) for visualization of FN and cell cytoskeleton, respectively. Cell nuclei were stained with DAPI. Cells were imaged with a Nikon Eclipse 80i epifluorescence microscope. Three biological replicas were analyzed.

For migration experiments, C2C12 myoblasts were cultured in starvation conditions (ultra-low serum conditions 1 % FBS) overnight for synchronization. 5 x 10^3^ C2C12 myoblasts cm^-2^ were seeded onto FN-functionalized glasses in the absence of FBS. The plate was disposed in the monitored stage of a Leica DMI6000 time-lapse microscope for recording cell movements. Images were recorded every 10 minutes for 24 h. Six migrating cells from two time-lapse videos per condition were analyzed using Fiji imaging software^102^.

### 4.16. Mutant FN

Cell adhesion assays were performed following the protocol described above. C2C12 myoblasts were seeded on glasses functionalized with 20 µg mL^-1^ of different mutant FN solutions (FN SYN, FN RGE, and FN ΔRGD) produced and purified as previously described^103^ in Dulbecco’s Phosphate Saline Buffer (DPBS) for 1 h at room temperature.

### 4.17. Image analysis

Focal adhesions were analyzed in Fiji imaging software^102^, as previously described^24^. Briefly, images were cropped to include only one cell per image, and background was subtracted using a sliding paraboloid and rolling ball radius of 50. Then, CLAHE was applied to enhance local contrast was using block size = 19, histogram bins = 256, and maximum slope = 6. The image underwent an exponential transformation to minimize background. In order to image brightness and contrast, automatic adjustments were performed, and then, a Log3D filter with sigmax = sigmay = 3 was applied. After inversion of the LUT of the image, the image was converted to a binary applying an automatic threshold. Then, a watershed algorithm was used to delete incorrectly clustered adhesions. Finally, FAs area and count per cell were quantified executing the “Analyze Particles” command (minimum 20 pixels to maximum infinity).

Cell area, aspect ratio and circularity were quantified in Fiji imaging software^102^. Briefly, a Gaussian blur filter (sigma = 2) was applied. Then, a default threshold was used to segment individual cells and quantify parameters of interest. The entire cell cytoskeleton was captured manually adjusting the threshold.

Actin retrograde flow at the cell edge speed was calculated by kymograph analysis in Fiji imaging software^102^. First, time-lapses were initially converted to 8-bit format, background was subtracted, contrast enhanced, and a Gaussian blur filter (sigma = 1.5) was applied. Then, a kymograph of line width 1 plotting displacement over time was produced using the Multi kymograph plugin. Actin retrograde flow speed was calculated using the bounding rectangle parameters and converting it from pixel units to nm units.

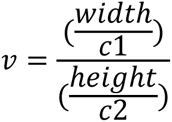

where c1 corresponds to the spatial conversion factor (px µm^-1^), and c2 is the temporal conversion factor (px s^-1^).

For matrix reorganization experiments, dark individual cell areas corresponding to reorganized FN deposited onto substrates by cells, were measured using Fiji imaging software^102^.

### 4.18. Statistics

All statistical analysis and graph plotting were performed in Prism v9 software (GraphPad). Data are presented as Mean ± SD. Normality tests were performed to determine whether to select parametric or non-parametric tests. Two-tailed unpaired student t-tests were employed when comparing two conditions. One-way ANOVA or two-way ANOVA, with post-hoc Tukey’s multiple comparisons test, were performed for comparisons involving more than two conditions with variations in one or two variables, respectively. Specific tests conducted for each analysis, sample size (n) and the meaning of the significance symbols are detailed in the respective figure captions. A p value at least smaller than 0.05 was chosen for significant comparisons. P values indicating significance: * ≤ 0.05, ** ≤ 0.01, *** ≤ 0.001, **** ≤ 0.0001.

## Acknowledgments

J.G-V acknowledges the funding from the European Union-NextGenerationEU and ARISTOS (European Union’s Horizon Europe research and innovation program under the MSCA grant agreement 101081334) programs. J.G-V acknowledges support by grant PID2022-137326OB-C22 funded by MCIN/AEI/10.13039/501100011033. P.R acknowledges support by grant PID2021-126012OB-I00 funded by MCIN/AEI/10.13039/501100011033 and by ERDF a way of making Europe, and by CIBER (CB06/01/1026). The authors acknowledge Mercedes Costell and Reinhard Fässler (Max Planck Institute of Biochemistry) for providing the mutant FN forms, and Pere Roca-Cusachs (IBEC) for the LifeAct-GFP plasmid. IBEC is member of CERCA Programme / Generalitat de Catalunya.

