## Supplementary Material for "NaBC1 Acts as a Mechanosensitive Co-regulator of Fibronectin-binding Integrins Adhesion and Myoblast Polarization"

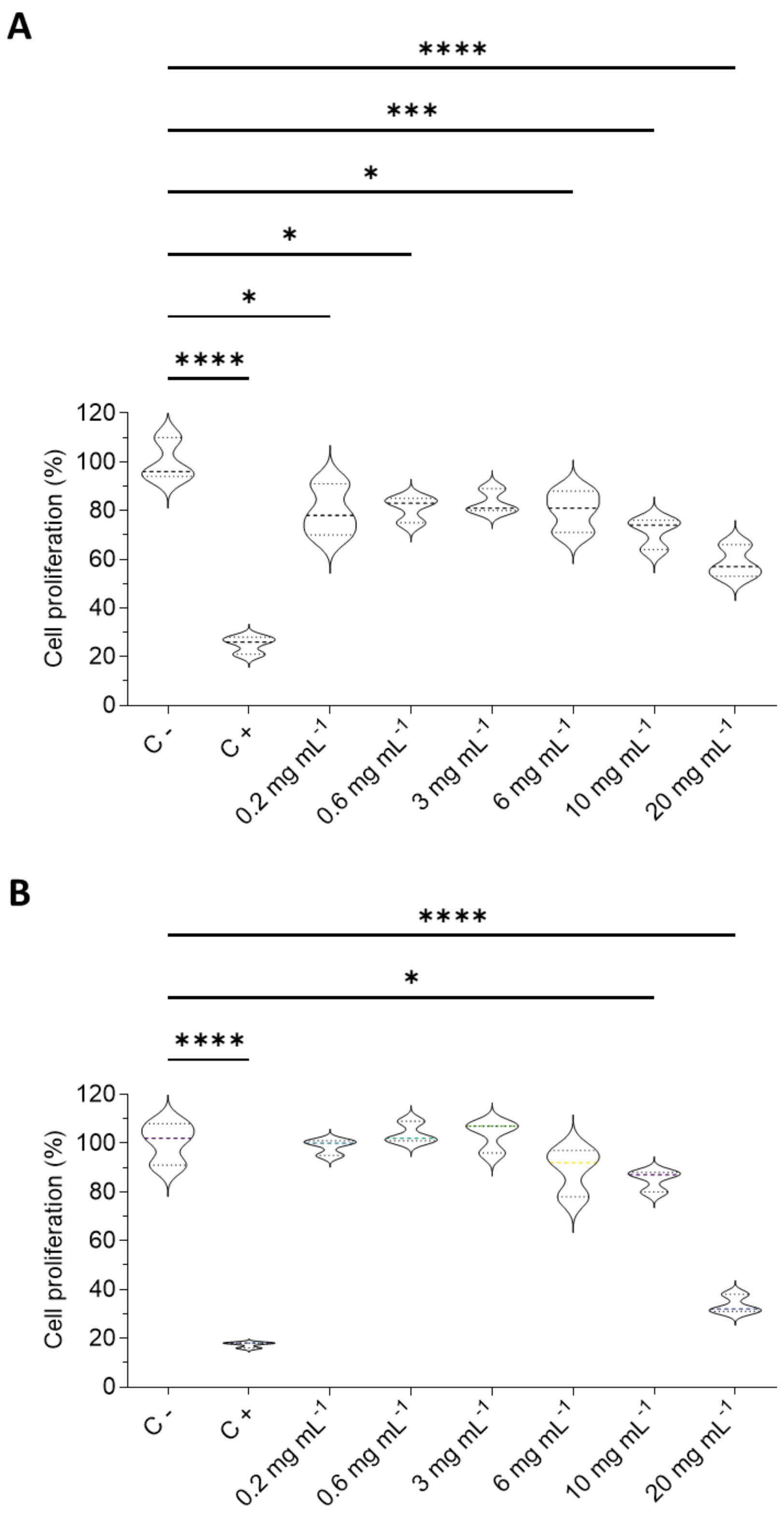


**Figure S1. Evaluation of borax (B) cytotoxicity.**

Quantified cell viability of C2C12 myoblasts seeded on FN-functionalized glasses and treated with B at the indicated concentrations (from 0.2 to 20 mg $mL^{-1}$) for 24 (A) and 48 h (B). n = 3 biological replicates with 3 technical replicates.

All data are presented as Mean ± SD. Statistical significance was determined using one-way ANOVA (Tukey's post hoc test) for multiple comparisons. *$p \leq 0.05$, ***$p \leq 0.001$, ****$p \leq 0.0001$

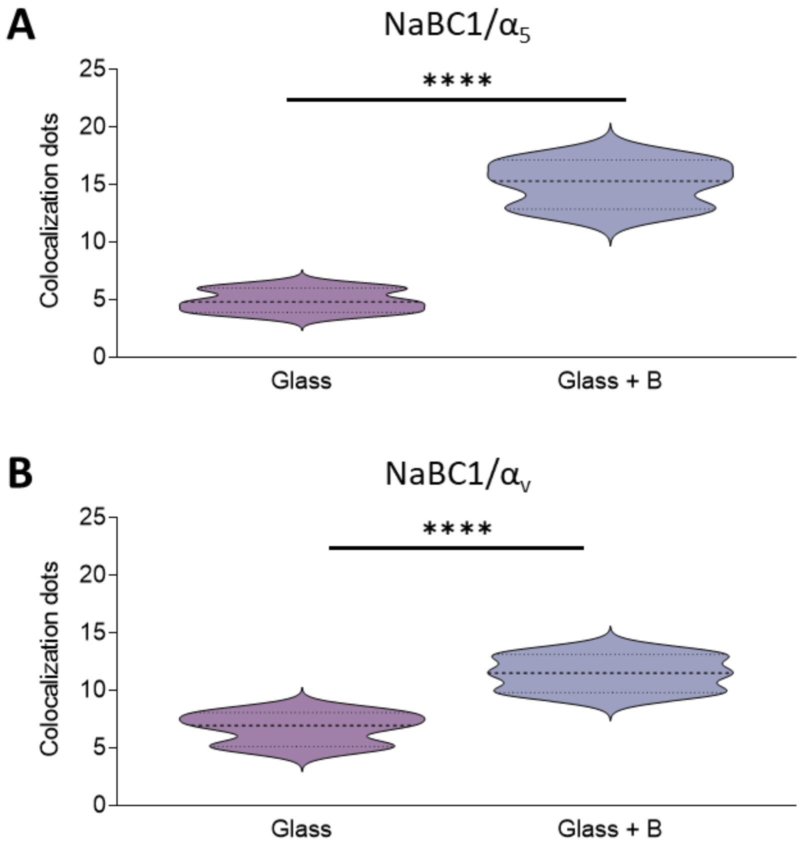


**Figure S2. Active-NaBC1 upregulates NaBC1-integrin colocalization.**

Quantified colocalization dots of NaBC1/$\alpha_5$ (A) and NaBC1/$\alpha_v$ (B) in C2C12 myoblasts seeded on FN-functionalized glasses and treated with 0.59 mM B for 0.5 h. n = 30 cells from 3 different biological replicates.

All data are presented as Mean ± SD. Statistical significance was determined using t-tests for pairwise comparisons. ****$p \leq 0.0001$

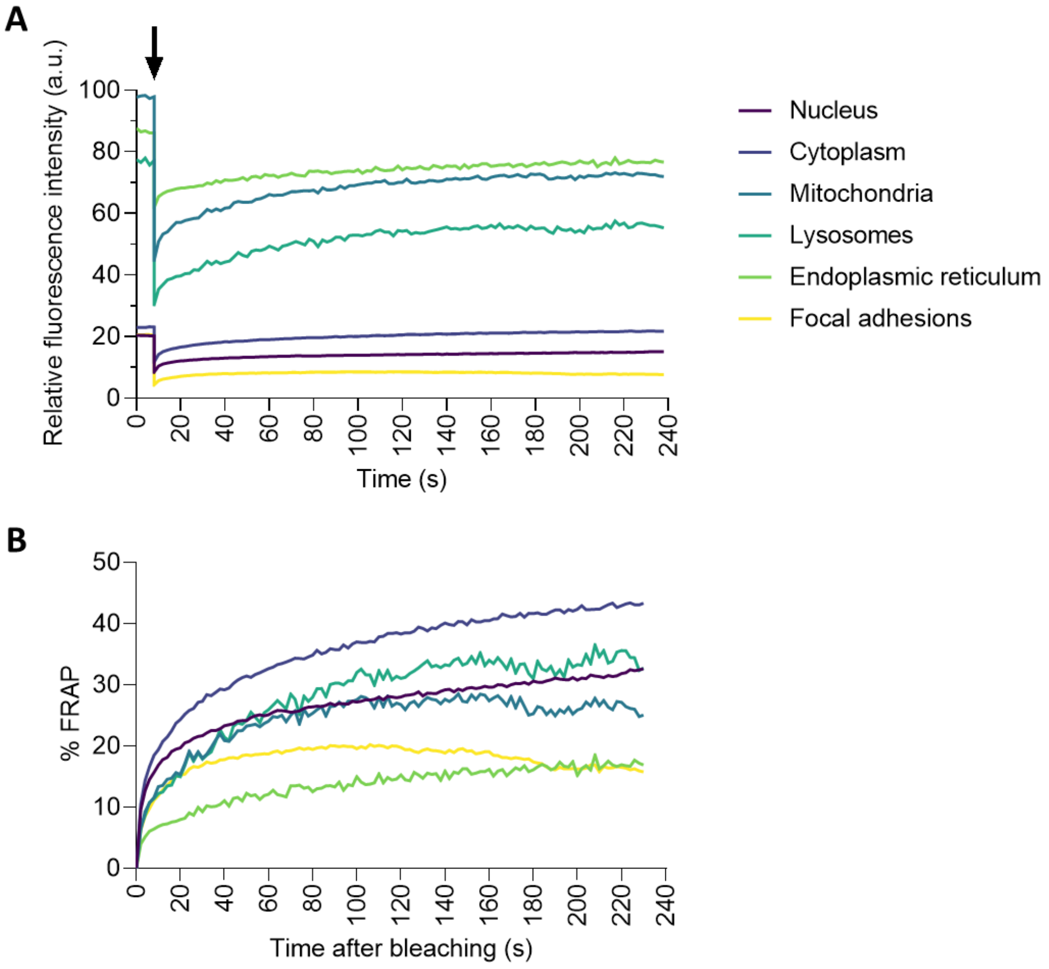


**Figure S3. FITC-B subcellular localization and dynamics.**

A: FRAP curves of FITC-labelled B fluorescence in C2C12 myoblasts seeded on FN-functionalized glasses and treated with FITC-labelled 0.59 mM B for 1 h. Arrow: bleaching time. n = 10 cells from 3 different biological replicates.

B: FRAP curves showing signal recovery after bleaching of fluorescence in C2C12 myoblasts, cultured as described in panel A. n = 10 cells from 3 different biological replicates. All data are presented as Mean.

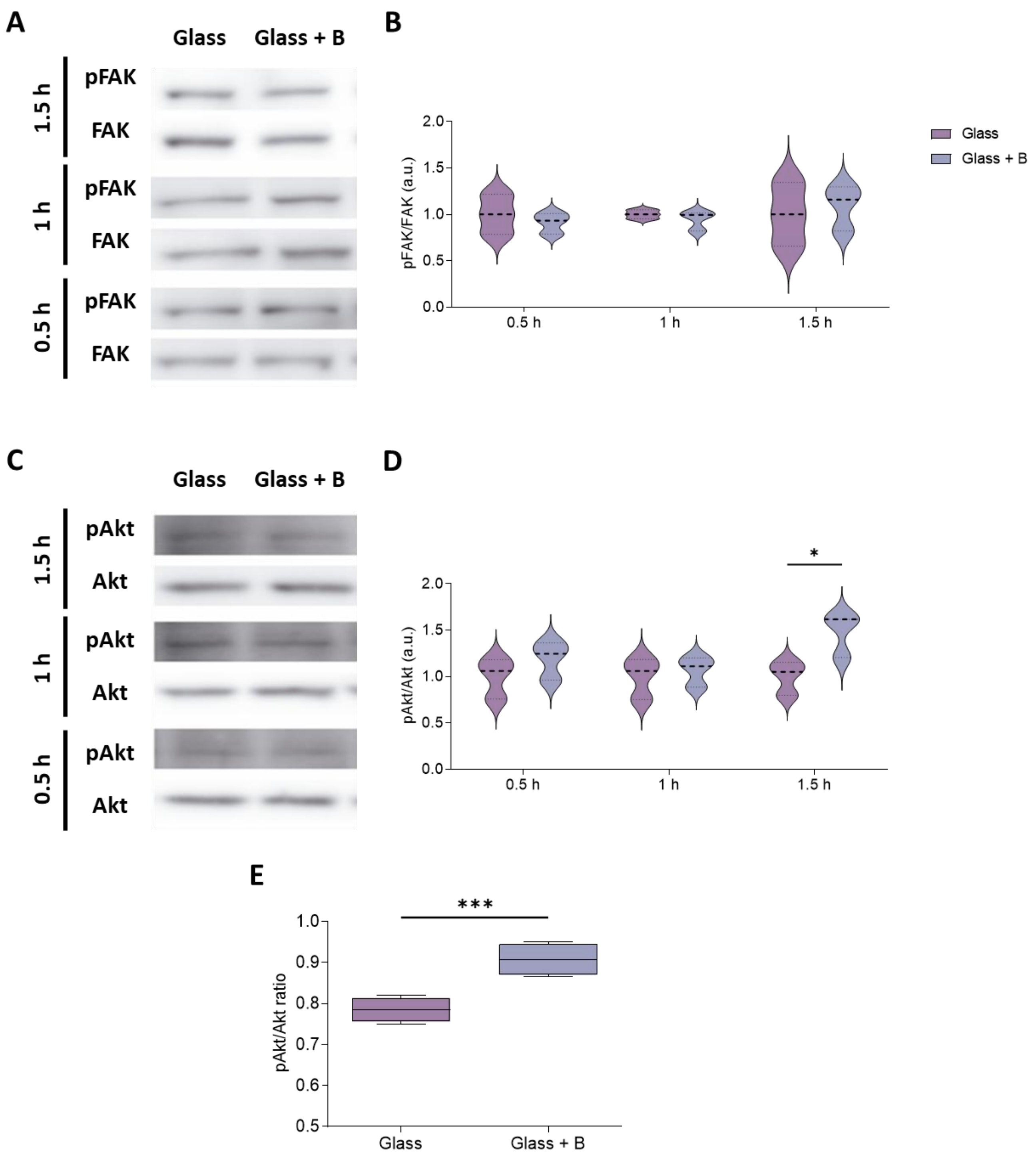


**Figure S4. The cluster NaBC1-FN-binding integrins triggers the Akt pathway.**

A: Western blot showing FAK and pFAK protein expression in C2C12 myoblasts seeded on FN-functionalized glasses and treated with 0.59 mM B for 0.5 h, 1 h and 1.5 h.

B: Quantified pFAK/FAK ratio protein expression.

C: Western blot showing Akt and pAkt protein expression in C2C12 myoblasts, cultured as described in panel A.

D: Quantified pAkt/Akt ratio protein expression.

E: Quantified pAkt/Akt ratio by In-Cell Western in C2C12 myoblasts seeded on FN-functionalized glasses and treated with 0.59 mM B.

n > 3 biological replicates with 3 technical replicates. All data are presented as Mean ± SD. Statistical significance was determined using two-way ANOVA (Tukey's post hoc test) for multiple comparisons (B and D) or t-tests for multiple or pairwise comparisons (E). $^*p \leq 0.05$, $^{****}p \leq 0.0001$

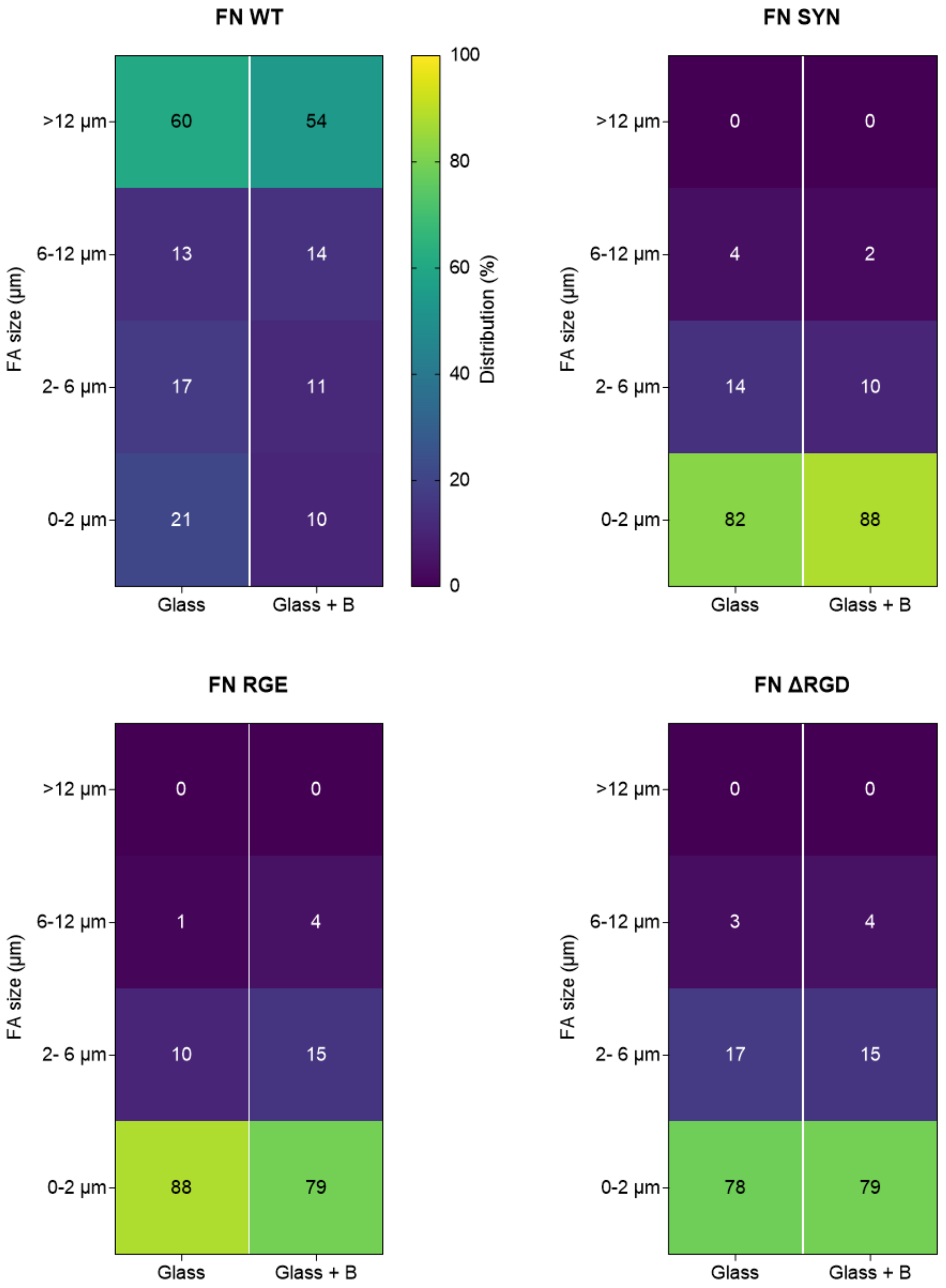


**Figure S5. The effect of active-NaBC1 on cell adhesion is dependent on RGD and synergy sites of fibronectin.**

Heat map of quantified FA size distribution in C2C12 myoblasts seeded on FN (WT or mutant FN forms)-functionalized glasses and treated with 0.59 mM B for 1.5 h. n > 50 imaged cells/condition from three different biological replicas.

All data are presented as Mean.
